# Spontaneous variability in gamma dynamics described by a linear harmonic oscillator driven by noise

**DOI:** 10.1101/793729

**Authors:** Georgios Spyropoulos, Jarrod Robert Dowdall, Marieke Louise Schölvinck, Conrado Arturo Bosman, Bruss Lima, Alina Peter, Irene Onorato, Johanna Klon-Lipok, Rasmus Roese, Sergio Neuenschwander, Wolf Singer, Martin Vinck, Pascal Fries

**Author notes:** These authors contributed equally to this work (Martin Vinck, Pascal Fries).

## Abstract

Circuits of excitatory and inhibitory neurons can generate rhythmic activity in the gamma frequency-range (30-80Hz). Individual gamma-cycles show spontaneous variability in amplitude and duration. The mechanisms underlying this variability are not fully understood. We recorded local-field-potentials (LFPs) and spikes from awake macaque V1, and developed a noise-robust method to detect gamma-cycle amplitudes and durations. Amplitudes and durations showed a weak but positive correlation. This correlation, and the joint amplitude-duration distribution, is well reproduced by a dampened harmonic oscillator driven by stochastic noise. We show that this model accurately fits LFP power spectra and is equivalent to a linear PING (Pyramidal Interneuron Network Gamma) circuit. The model recapitulates two additional features of V1 gamma: (1) Amplitude-duration correlations decrease with oscillation strength; (2) Amplitudes and durations exhibit strong and weak autocorrelations, respectively, depending on oscillation strength. Finally, longer gamma-cycles are associated with stronger spike-synchrony, but lower spike-rates in both (putative) excitatory and inhibitory neurons. In sum, V1 gamma-dynamics are well described by the simplest possible model of gamma: A linear harmonic oscillator driven by noise.

---

The brain consists of different kinds of cell types which have unique properties, and are commonly divided into inhibitory (I) and excitatory (E) neurons. Interactions among I and E neurons can generate collective rhythmic activity in different frequency bands. One of the “faster” rhythms that neocortical circuits can generate is the gamma rhythm (30-80Hz), whose function has been heavily debated in the literature ^1-20^. This rhythm can be observed at many scales, from the macro/meso-scale (MEG, EEG, ECoG, LFP), to the microscale (synaptic currents and spiking activity) ^10, 21 22^. It is however unknown how the properties of collective neuronal gamma synchronization can arise from the interactions between its microscopic constituents ^21, 23^.

Observations of macro/meso-scopic gamma dynamics have revealed substantial variability in the amplitude and frequency of gamma oscillations as a function of time, but also cortical space ^7, 16, 24-29^. In particular, gamma oscillations are not well approximated by sinusoids ^7^, despite the fact that they are often depicted as such. Rather, they show major fluctuations in their amplitude over time, sometimes described as “bursts”; as well as their frequency, giving rise to the broad-band spectral nature of gamma. These fluctuations likely reflect the properties of the underlying E-I circuit and the way it responds to changes in input drive, and they impose constraints on the possible functional roles of gamma ^8, 9, 13, 15, 16, 18, 28, 30^. A previous study in rodent hippocampus ^31^ has suggested that cycle-by-cycle fluctuations in amplitude and duration (i.e. the inverse of frequency) are explained by two components: (1) cycle-by-cycle fluctuations in synaptic excitation; and (2) balanced, bidirectional interactions between E and I neurons, consistent with the PING (Pyramidal Interneuronal Network Gamma) model of the gamma rhythm ^5, 10, 32-37^. This model holds that the occurrence of a strong bout of synaptic excitation is balanced by high-amplitude, long-lasting inhibition. As predicted from this model, this study reported that gamma-cycle amplitude and duration are strongly correlated (r = 0.61) in rodent hippocampus ^31^.

The starting point of the present study was to see whether this regularity generalizes to other cortical circuits, in particular to awake primate visual cortex, another system where gamma oscillations have been extensively studied. It remains unclear how the mechanisms of gamma in visual cortex compare to hippocampus. It appears that E-I mechanisms of gamma in higher visual areas (V4) might be comparable to hippocampus ^37^, although there is evidence that they are substantially different in primary visual cortex (V1) ^38^. Furthermore, the dependence of V1/V2 gamma on stimulus contrast suggests that increases in synaptic excitation lead to *increases* rather than *decreases* in the frequency of V1/V2 gamma ^25, 39, 40^. It is unknown, however, what the relationship is between *spontaneous* fluctuations in gamma-cycle amplitude and duration in area V1.

## RESULTS

### Recordings and Task

We recorded LFPs and spiking activity from the primary visual cortex (V1) of several awake macaque monkeys (see Methods). Monkeys performed a fixation task, while drifting gratings or uniform colored surfaces were presented. Fig. 1a shows an example trial of broad-band LFP recorded during the presentation of a full-screen drifting grating. The trial-average spectra of absolute power (Fig. 1b) and of the power-change relative to pre-stimulus baseline (Fig. 1c) reveal strong visually-induced gamma oscillations. The time-frequency analysis (Fig. 1d) shows that this induced gamma rhythm is sustained for the duration of the visual-stimulation period. Fig. 1f-i shows similar results for visual stimulation with a colored surface ^41, 42^.

**Figure 1.**
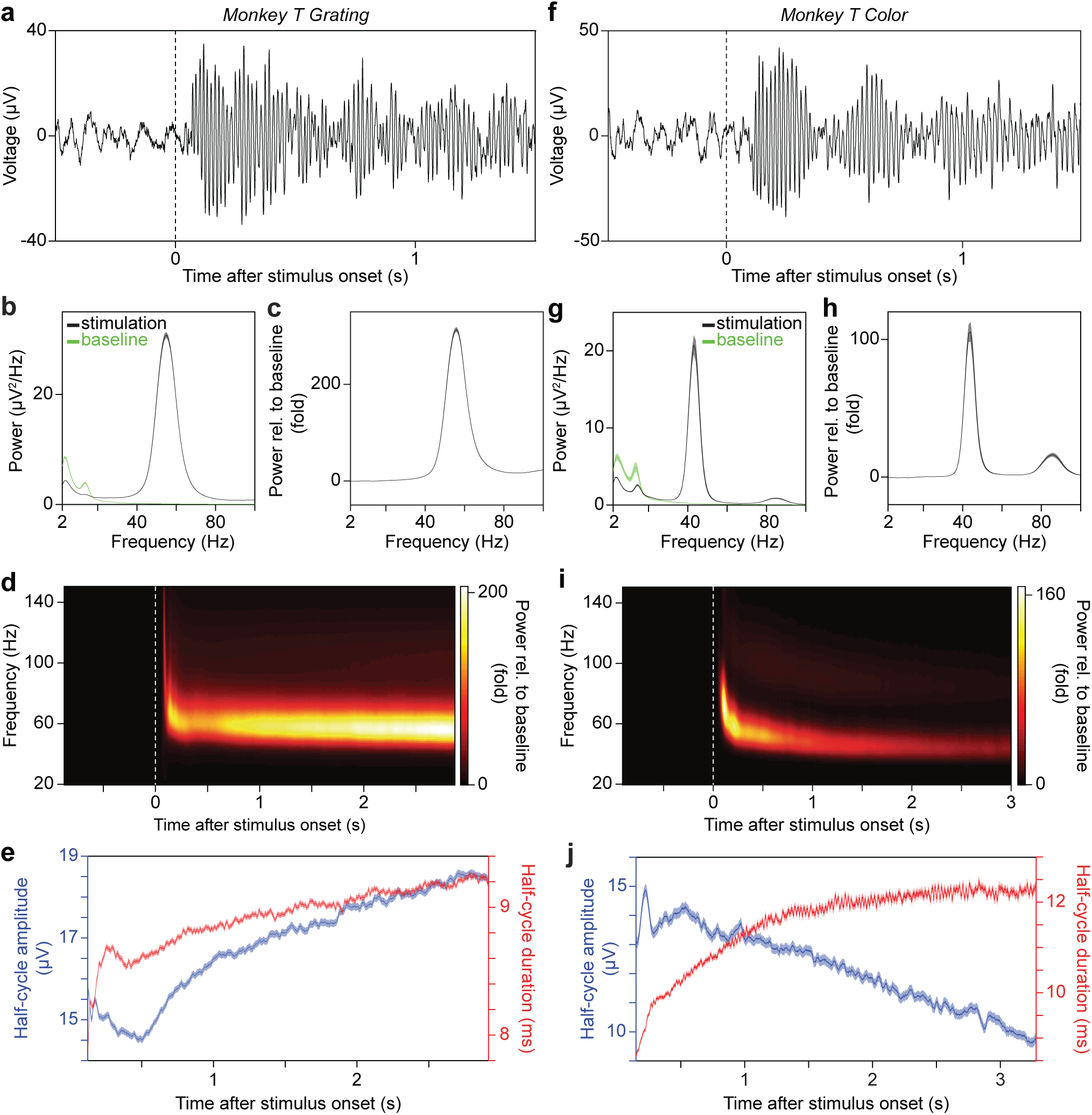
Gamma dynamics in awake macaque V1 during visual stimulation. (**a**) Raw LFP trace from one representative recording site from area V1 in monkey T before and during the presentation of a full-screen drifting grating. (**b**,**c**) Raw power (**b**) and power change relative to baseline (**c**), averaged across all selected recording sites from V1 in monkey T. The green and black traces in **b** correspond to the pre-stimulus baseline period and stimulation period respectively. The error regions show 2 standard errors of the mean (S.E.M.) based on a bootstrap procedure across trials (1000 bootstraps). (**d**) Power change relative to baseline, as function of frequency and time relative to stimulus onset, averaged over all selected V1 recording sites in monkey T before and during the presentation of a full-screen drifting grating. Note the changes in gamma amplitude and frequency with time after stimulus onset. (**e**) Time course of gamma-half-cycle amplitude (blue) and duration (red), averaged over all selected V1 recording sites in monkey T during the presentation of a full-screen drifting grating. The error regions show ±2 SEM based on a bootstrap procedure. Only the stimulation period is shown, because only very few gamma cycles of very low amplitude were detected before stimulus onset. (**f**-**j**) Same as **a**-**e**, but for the presentation of a full-screen uniform color surface. (**a**,**d**,**f**,**i**) Dashed lines indicate stimulus onset.

### The correlation between gamma cycle amplitude and duration

A previous study has examined correlations between the amplitude and duration of individual gamma cycles in the CA3 field of the rat hippocampus ^31^. This study found a strongly positive (r = 0.61) correlation between amplitudes and durations, both in vivo and in vitro. We wondered whether a similarly strong correlation exists in monkey V1. We therefore used the same analysis method as previously used for the rat hippocampus. This method is based on (1) band-pass filtering LFP signals, (2) detecting periods of high-amplitude gamma activity, and (3) detecting empirical peaks and troughs in the filtered signal (Fig. 2a,b; see Methods). Using this method, we found a relatively strong positive (r = 0.361) correlation between the amplitude and duration of individual gamma cycles in the visual stimulation period (Fig. 2c). By contrast, correlations between the amplitude of a given cycle and the duration of either the preceding or succeeding cycle were not significant (Fig. 2c).

**Figure 2.**
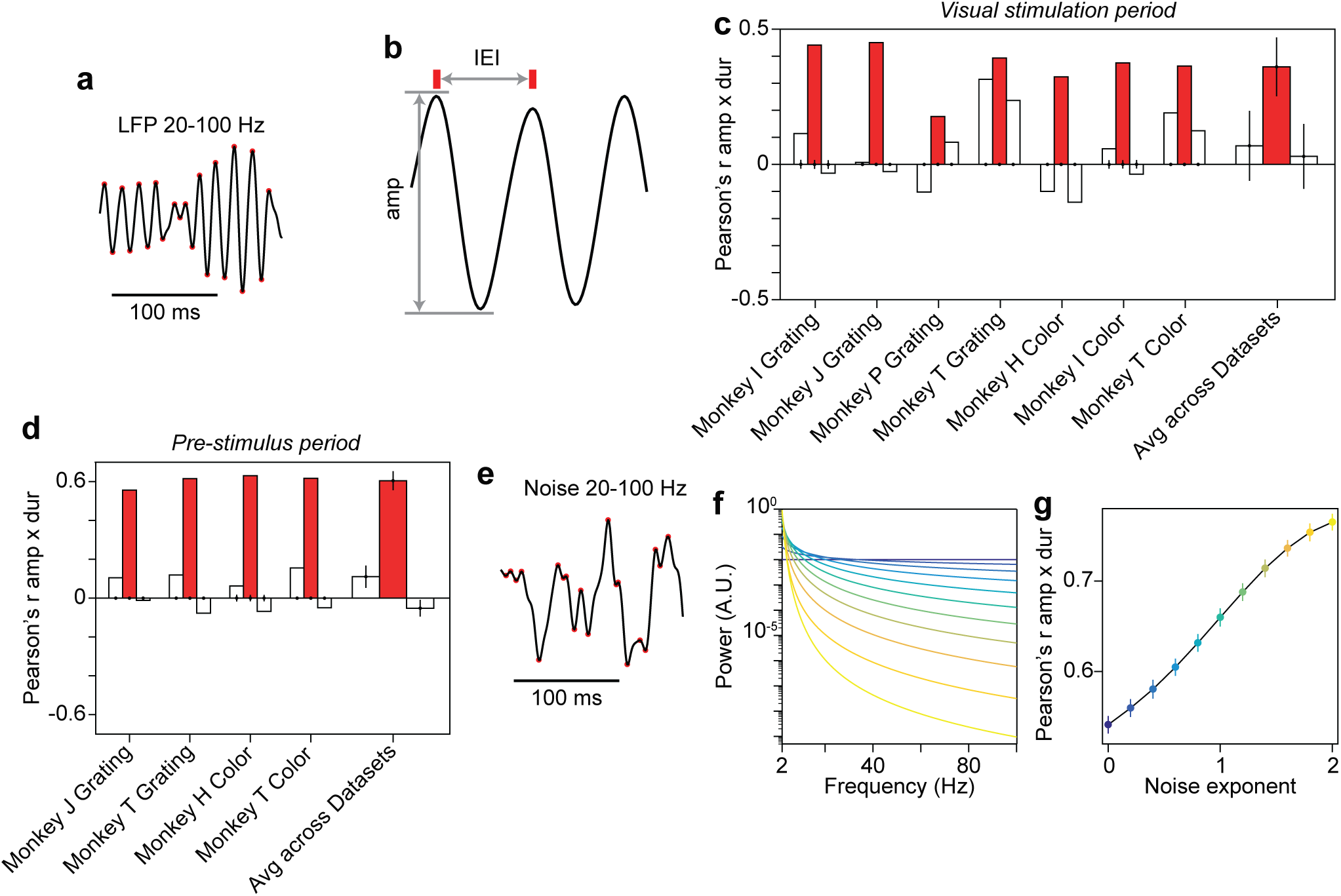
Estimation of correlation between gamma-cycle amplitude and duration can be influenced by noise. (**a**) LFP trace filtered in the gamma range (20-100 Hz). Red dots indicate local maxima and minima. (**b**) Segment of the trace in **a** demonstrating the definition of gamma-cycle amplitude and inter-event interval, i.e. gamma-cycle duration. (**c**) For each dataset listed on the x-axis, the three bars show the correlation between gamma-cycle amplitudes and the durations of the same gamma cycle (center, red), the previous gamma cycle (left, white) and the next gamma cycle (right, white). On the right, this is shown for the average across all datasets. This was calculated for the visual stimulation period. Amplitude and duration values were extracted as in ^31^, including the filtering illustrated in (**a, b**); note that the employed subtraction of a boxcar-smoothed signal amounts to a high-pass-filtering at 20 Hz. For each dataset, a null distribution was produced by randomizing the order of duration values across trials, and the resulting means and 99.9% confidence intervals are shown as dots and vertical lines. For the average across datasets, shown on the right, we performed a t-test and show the resulting confidence intervals as vertical lines on the observed mean (red bar: p<5*10^−5^, white bars for preceding cycle: p=0.28, white bars for succeeding cycle: p=0.56). In addition, we performed a two-sided non-parametric permutation test (red bar: p<0.05; white bars: p>0.05). (**d**) Same as **c** but for the pre-stimulus baseline (averages across datasets: red bar: p=4.51*10^−5^, t-test across datasets; white bars p=0.011 and p=0.038, respectively for preceding and succeeding cycles, t-test across datasets) (**e**) Example synthetic colored noise trace filtered in the gamma range (20-100 Hz). Red dots indicate local maxima and minima. (**f**) Power spectra of synthetic colored noise signals with a spectral shape of 1/f^n^, with n assuming values from 0 (dark blue) to 2 (bright yellow). (**g**) Correlation of the amplitude and duration of individual deflections in synthetic colored noise signals. Dots and vertical lines indicate means ±2 SEM produced by a bootstrap procedure (1000 bootstraps). The color conventions are the same as in **f**.

We expected that this result would be specific to the visual stimulation period, in which gamma oscillations were prominent, but that it would not hold true for the pre-stimulus period, in which there was no visible gamma peak in the LFP power spectrum (Fig. 1b,g). Nonetheless, for the pre-stimulus period, the algorithm detailed above detected a substantial amount of gamma epochs. Surprisingly, we observed even stronger correlations between gamma-cycle amplitudes and durations for the pre-stimulus (r = 0.605) compared to the stimulus period (Fig. 2d).

This prompted us to investigate whether the same algorithm would also detect a positive correlation between gamma-cycle amplitudes and durations for synthetic 1/f^n^ noise signals (Fig. 2e,f). This was indeed the case (Fig. 2g). Thus, noisy fluctuations in a signal without rhythmic components can give rise to a strong positive correlation between the amplitudes and durations of detected “gamma cycles”. The presence of a positive correlation between amplitudes and durations can be made intuitive by considering a random walk process: In such a process, the magnitudes of successive steps (i.e. increments or decrements) are independent of each other, with zero mean. In this case, a successive series of positive increments typically results in a “cycle” with a high amplitude and a long duration. By contrast, a rapid reversal typically results in a low-amplitude “cycle” with a short duration. Together, these findings indicate that the positive correlation between gamma-cycle amplitude and duration in the stimulus period may have been due to noisy background fluctuations.

These results prompted us to develop a method that (1) avoided band-pass filtering in a narrow frequency-range; and (2) ensured that gamma peaks and troughs were not detected due to noisy fluctuations, but reflected a rhythmic process (Fig. 3a-d; see Methods). To obtain estimates of gamma-cycle amplitudes and durations with a high temporal resolution, we measured them in periods of “half-cycles” (i.e. peak-to-trough or trough-or-peak). (For the rest of the text, we will be referring to the amplitudes and durations of individual gamma half-cycles as “gamma-cycle amplitudes” and “gamma-cycle durations”, and will mention explicitly when we measure them in full rather than half cycles). In contrast to the method used for Fig. 2, we found that our method detected very few gamma cycles in the pre-stimulus period (Fig. 3a-c). Because of this, a correlation between gamma-cycle amplitude and duration could not be reliably computed for this period. To further examine the noise-robustness of our method, we simulated an AR(2) (2^nd^ order auto-regressive) process that had a positive correlation between gamma-cycle amplitudes and durations in the absence of noise. We then added 1/f^2^ background noise of different intensities (see Methods). We found that our method did not yield spurious correlations due to the inclusion of noise; instead it failed to detect any gamma cycles for higher noise-levels (red line in Fig. 3e). By contrast, the method used for Figure 2 produced higher correlations as the noise-level increased (Fig. 3e).

**Figure 3.**
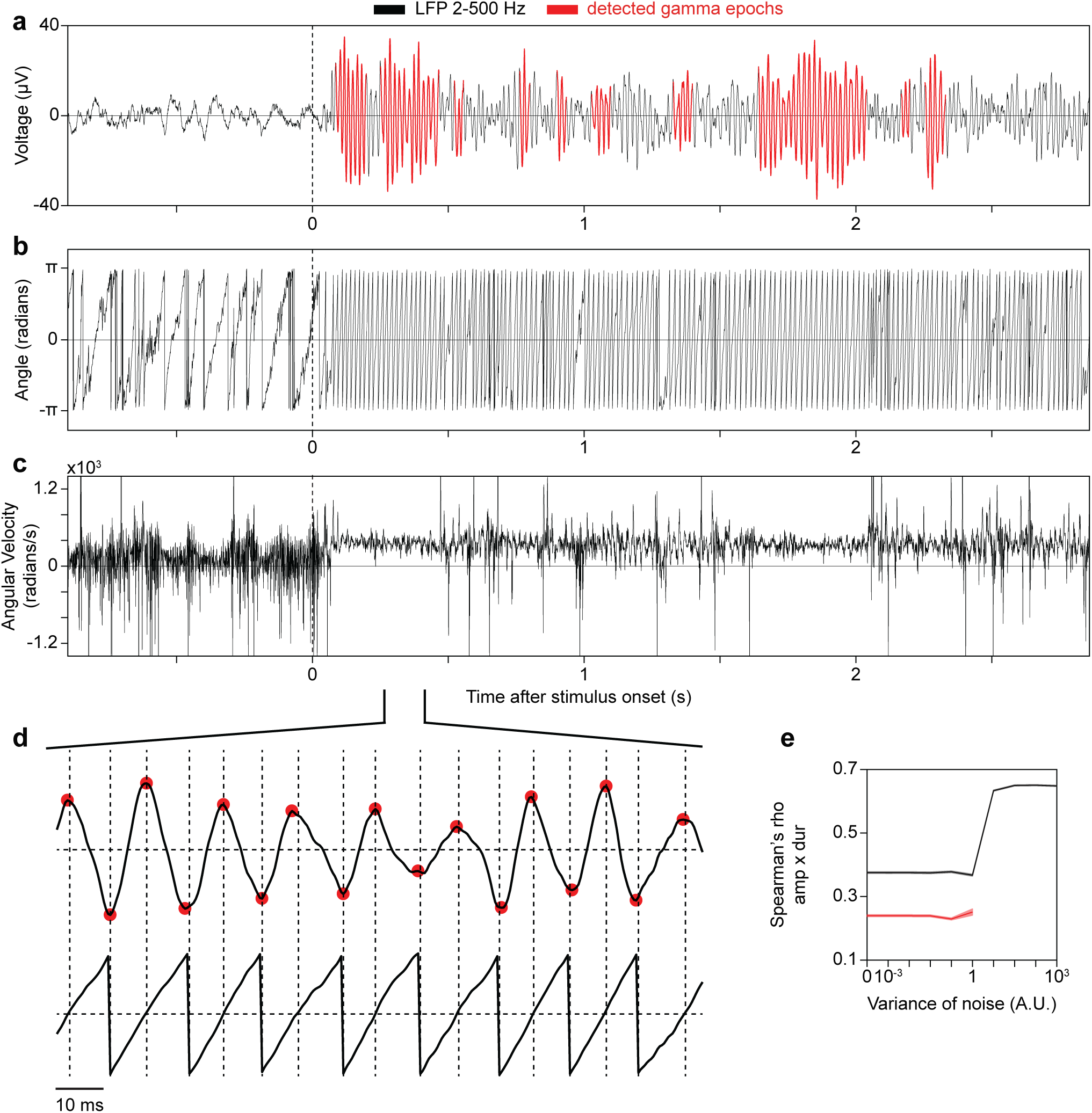
Illustration of a method for the selection of gamma-oscillatory epochs. (**a**) LFP trace displayed in **Fig. 1a**, with regions presented in red corresponding to gamma epochs passing the criterion for stationarity. (**b**) Phase of the analytic signal based on the Hilbert transform of the trace shown in **a**. (**c**) Angular velocity of **a**. Note periods of positive and relatively stable angular velocity, corresponding to oscillatory gamma epochs in the original LFP. (**a**-**c**) Dashed lines indicate stimulus onset. (**d**) Magnification of the designated section of the LFP trace and its phase. Red dots indicate detected LFP peaks and troughs. Vertical dashed lines designate negative-to-positive and positive-to-negative zero crossings of the phase of the analytic signal, whereas horizontal dashed lines designate 0. (**e**) The correlation between the amplitude of a gamma half-cycle and the duration of the same gamma half cycle for different additive noise levels, computed with the method used in **Fig. 2** (black), and the method described in this figure (red). The error regions show ±2 SEM based on a bootstrap procedure.

Using this new method, we then detected gamma-cycle amplitudes and durations for all trials and available time-points, separately for each recording site and stimulus condition. Because we were interested in spontaneous variability, we further ensured that correlations between gamma-cycle amplitudes and durations could not arise due to the time courses of amplitude and frequency after stimulus onset (Fig. 1e,j). We achieved this by computing correlations across trials, separately for each available post-stimulus time-point and then averaging the correlations over time-points (see Methods). With this approach, we found that the amplitudes and durations of individual gamma half-cycles were positively correlated in all tested datasets (Fig. 4a). The magnitude of these correlations was, on average, substantially lower (rho = 0.199) than the one observed with the previously employed method (compare Figs 2c and 4a). In addition, we computed the correlation between the amplitude of a given half-cycle and the duration of the previous or the subsequent half-cycles, and this did not result in a consistent pattern of correlations across datasets (white bars in Fig. 4a). Similar results were obtained for full rather than half cycles, with significant correlations only for the same cycle comparison, but not for the preceding and succeeding cycle (Supplementary Fig. 1a). Thus, amplitude and duration were weakly but positively correlated across individual cycles of awake monkey V1 gamma.

**Figure 4.**
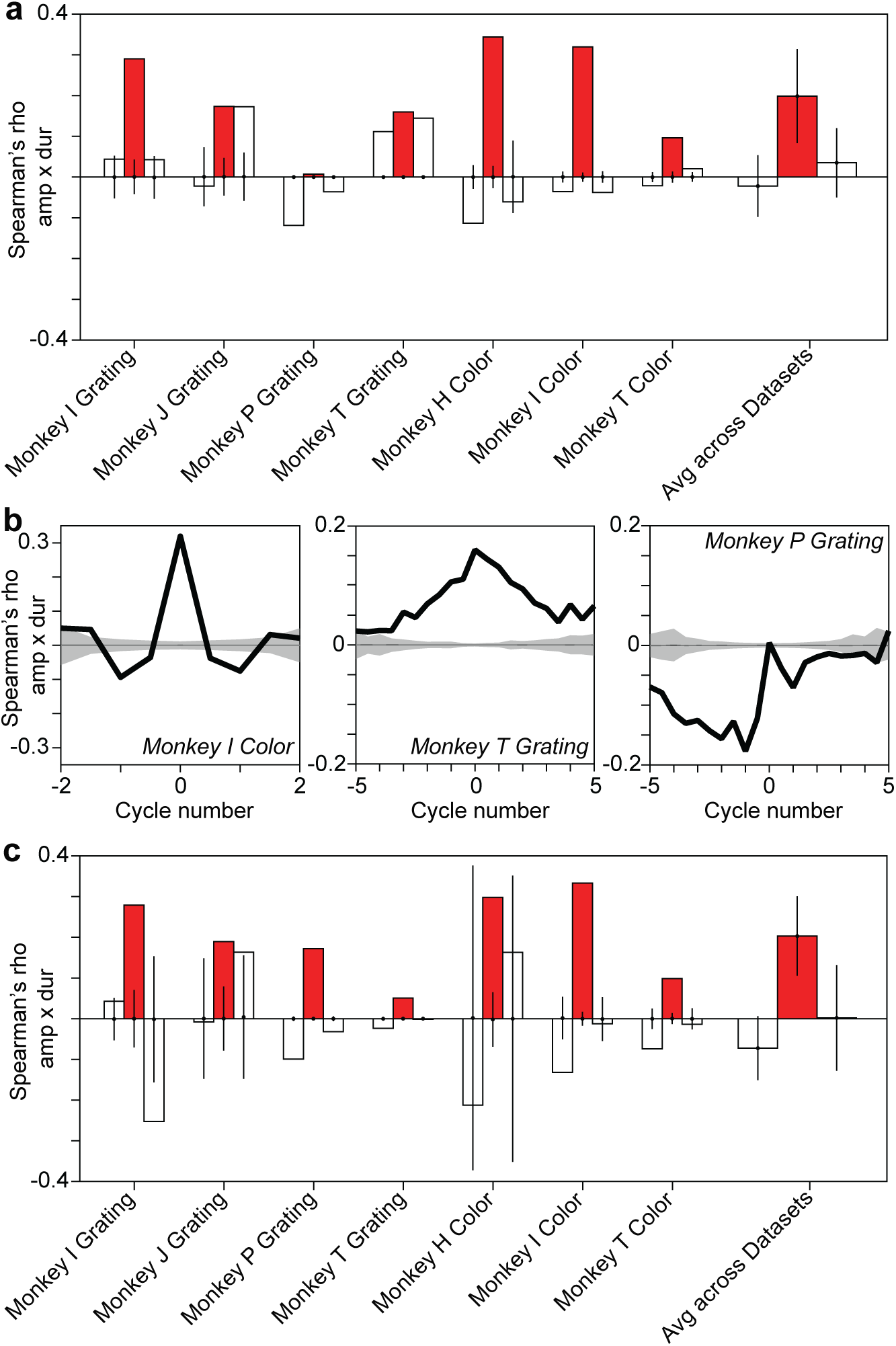
Gamma-half-cycle amplitudes and durations are positively correlated in gamma-oscillatory epochs. (**a**) For each dataset listed on the x-axis, the three bars show the correlation between the amplitude of a gamma half cycle and the duration of the same (center, red), previous (left, white) and next (right, white) gamma half cycle. On the right, this is shown for the average across all datasets. This was calculated for each time point across trials and averaged across time points for gamma-oscillatory epochs. The data used correspond to the period during the presentation of the visual stimulus. For individual datasets, a null distribution was produced by randomizing the order of duration values across trials, and the resulting means and 99.9% confidence intervals are shown as dots and vertical lines. For the average across datasets, shown on the right, we performed a t-test and show the resulting confidence intervals as vertical lines on the observed mean (red bar: p<6*10^−3^, white bars for preceding cycle: p=0.5, white bars for succeeding cycle: p=0.35). In addition we performed a two-sided non-parametric permutation test (red bar: p<0.05; white bars: p>0.05). (**b**) Correlation between the amplitude of a gamma half-cycle and the duration of gamma half-cycles before and after it for 3 different datasets. Note that in monkey I, this is limited to ±2 cycles, because the signal-to-noise ratio was lower, resulting in shorter gamma-oscillatory epochs. Importantly, all three example datasets show a central peak, despite the fact that they show different longer-term correlations. The gray lines and gray-shaded areas depict the means and 99.9% confidence regions, after randomizing the order of duration values across trials. (**c**) Same as **a**, but showing the correlations between residuals of the regression across adjacent amplitude triplets and the residuals of the regression across adjacent duration triplets (red bar: p=23*10^−4^, t-test across datasets; p<0.05, permutation test for individual datasets; white bars p=0.066 and p=0.97, respectively for preceding and succeeding cycles, t-test across datasets; p>0.05, permutation test for individual datasets).

### The influence of slow dynamics and microsaccades

We wondered whether the observed correlation between gamma-cycle amplitudes and durations may have resulted from correlated changes in amplitudes and durations at relatively slow time scales, e.g. due to drifts or slow oscillations in the monkey’s state, or stimulus repetition effects ^42-44^. In order to control for the potential influence of such changes, we computed the correlation between the amplitude of a given half-cycle and the duration of multiple preceding and succeeding half-cycles (Fig. 4b). Some datasets showed dynamics on the temporal scale of few half-cycles (Fig. 4b, left), and others on the scale of multiple half-cycles (Fig. 4b, middle and right). For example, the right panel in Fig. 4b shows a long-lasting, negative trend punctuated by a small positive value for the instantaneous correlation. By contrast, the middle panel shows a positive trend peaking at zero lag. These trends may have contributed to the observed correlation between gamma-cycle amplitude and duration. We therefore removed the influence of slower dynamics through a linear regression analysis (see Methods). In this analysis, we first regressed out linear predictions of gamma-cycle amplitude and of duration from previous and succeeding cycles, and repeated the analysis on the regression residuals (see Methods). We found that the resulting correlation was comparable to the correlation between raw amplitude and duration (compare Fig. 4a and 4c). Similar results were obtained for full cycles (Supplementary Fig. 1b). Together, these findings indicate that the positive correlation between gamma-cycle amplitudes and durations was not due to within- or across-trial trends on a longer timescale. Further analyses also suggest that the correlation between gamma-cycle amplitudes and durations was not due to transient changes in amplitudes and durations following microsaccades (Supplementary Fig. 2; see Methods).

To further understand the contribution of non-stationarities to the correlation between gamma-cycle amplitudes and durations, we fitted an autoregressive (AR) model to the LFP data. An AR model captures the variance and auto-correlation of the LFP, and can then be used to generate a stationary surrogate time-series, (by stationary we mean that the underlying statistics of the signal do not change over time). Supplementary Fig. 3a-d illustrates this for the dataset used for Fig. 1a-e. We find that the AR model accurately captured the power spectrum (Supplementary Fig. 3b), but did not replicate slower dynamics in gamma-cycle amplitudes or durations (Supplementary Fig. 3c,d; compare to Fig. 1d,e). In the surrogate data generated by the AR model, we then analyzed the correlations between gamma-cycle amplitudes and durations, and found consistently positive correlations of similar average strength as in the original data (Supplementary Fig. 3e). Again, similar results were obtained for full rather than half cycles (Supplementary Fig. 1d). These results further support the notion that the observed correlations in the LFP data were not due to co-fluctuations or non-stationarities on a slower time scale.

### The cycle-based amplitude spectrum and the rate of incidence of cycle-durations

In the V1 LFP data, we observed a small but positive correlation between gamma-cycle amplitudes and durations. This correlation, however, does not necessarily imply a monotonic or linear relationship between gamma-cycle amplitudes and durations, as was reported by Atallah and Scanziani (2009). We thus examined the joint distribution of gamma-cycle amplitudes and durations in more detail. To this end, we first computed the average half-cycle amplitude for each possible half-cycle duration (Fig. 5a, see Methods); we refer to this as the cycle-based amplitude spectrum (CBAS). To minimize the possible influence of stimulus-locked trends in gamma amplitude and frequency, we used only the final 250 ms of visual stimulation. To average CBASs across monkeys, we first converted half-cycle duration values to frequency values (in Hz). We then aligned the CBASs to the “gamma peak frequency”, that is the frequency at which the Fourier-based power spectrum (FBPS) reached a maximum. In the CBAS, we found that the relationship between frequency and amplitude was non-monotonic: The amplitude was greatest at a frequency that was slightly lower than the peak gamma frequency, and showed a decline towards higher gamma frequencies. In contrast to the CBAS, we observed that the FBPS was approximately symmetric (Fig. 5a). Thus, FBPS had a different shape and dependence on frequency than the CBAS. We further wondered how often different gamma-cycle durations tended to occur. We therefore computed the cycle-frequency (i.e. inverse of gamma-cycle duration) distribution. We found that the cycle-frequency distribution was approximately symmetric, and closely matched the FBPS. Specifically, we found that the most prevalent half-cycle frequency lied within one Hertz of the peak gamma-frequency derived from the Fourier-based power spectrum (Fig. 5A and Supplementary Fig. 4a).

**Figure 5.**
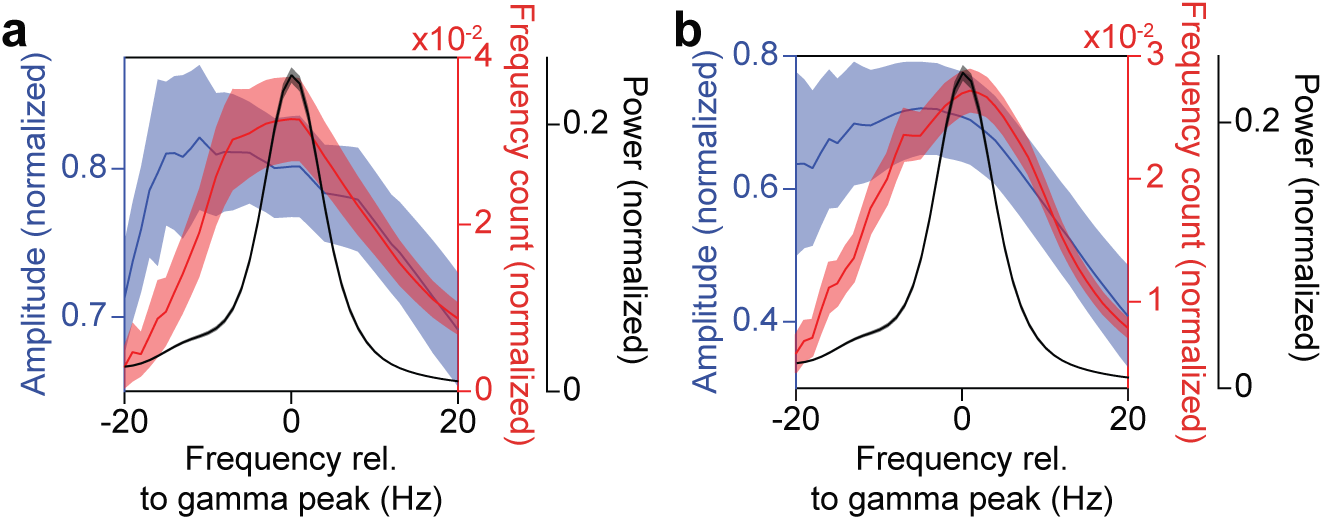
Cycle-based amplitude-spectra and cycle-frequency distributions. (**a**) The x-axis shows duration expressed as its inverse, namely frequency, and after aligning to the gamma peak in the raw power spectrum (black trace). The blue curve shows the gamma-half-cycle amplitudes as a function of their duration. The red curve shows the count of detected gamma half-cycles as a function of their duration. These analyses were based on the broadband signal from the last 250 ms of stimulation (see Methods). Error regions show ±2 SEM based on a bootstrap procedure. (**b**) Same as **a**, but for gamma epochs detected on the filtered LFP.

We wondered whether the observed dependency of gamma-cycle amplitude on cycle-frequency may have been due to a ceiling effect, because in our analysis we selected those broad-band LFP segments for which gamma rhythms were relatively strong. This selection circumvented several methodological problems, as discussed above and in the Methods section. Yet, it may have limited the generalizability of our findings. To address this issue, we re-analyzed the data after band-pass filtering the LFP in the gamma-frequency range (20-100 Hz). This modification in our approach substantially increased our sensitivity in detecting gamma episodes. The distributions of cycle-frequency and amplitude that we obtained after band-pass filtering were, nevertheless, highly similar to the ones calculated on the broad-band signal (Fig. 5b and Supplementary Fig. 4b).

### Relationship of gamma frequency with spiking

We further wondered how the spontaneous dynamics of gamma oscillations related to the activation and phase-locking of excitatory and inhibitory neurons. The more complex PING (Pyramidal Interneuron Network Gamma) model of Atallah and Scanziani (2009), discussed above, predicts that higher-amplitude gamma cycles are initiated by a stronger bout of excitatory spiking. These excitatory bouts should then give rise to longer-lasting inhibition, resulting in longer gamma cycles ^31^.

In order to assess if this prediction holds true for awake macaque V1, we analyzed multi-unit (MUA) activity (see Methods). We first computed the normalized spike count (number of spikes per cycle) (Fig. 6a) as a function of the gamma-cycle frequency (the inverse of gamma-cycle duration; see Methods). The normalized spike count was negatively correlated with gamma-cycle frequency (Fig. 6d). This may be a trivial result, because the spike count may simply reflect the product of firing rate and gamma-cycle duration. To correct for this, we divided the spike count by the duration, yielding the firing rate (spikes/sec). We observed that firing rates were positively correlated with gamma-cycle frequency. We wondered if the same result holds true for different excitatory and inhibitory cell classes. For this reason, we classified single units into three classes that were previously identified by Onorato et al. (2020) in the same dataset: NW-Burst, NW-Nonburst (NW: Narrow waveform) and BW (Broad Waveform) units. Previous studies suggest that NW-Burst and BW units correspond to putative pyramidal cells, whereas NW-Nonburst neurons correspond to putative fast-spiking interneurons ^37, 45, 46^. We found that firing rates were positively correlated with frequency for all three classes, similar to the MUA (Fig. 6d).

**Figure 6.**
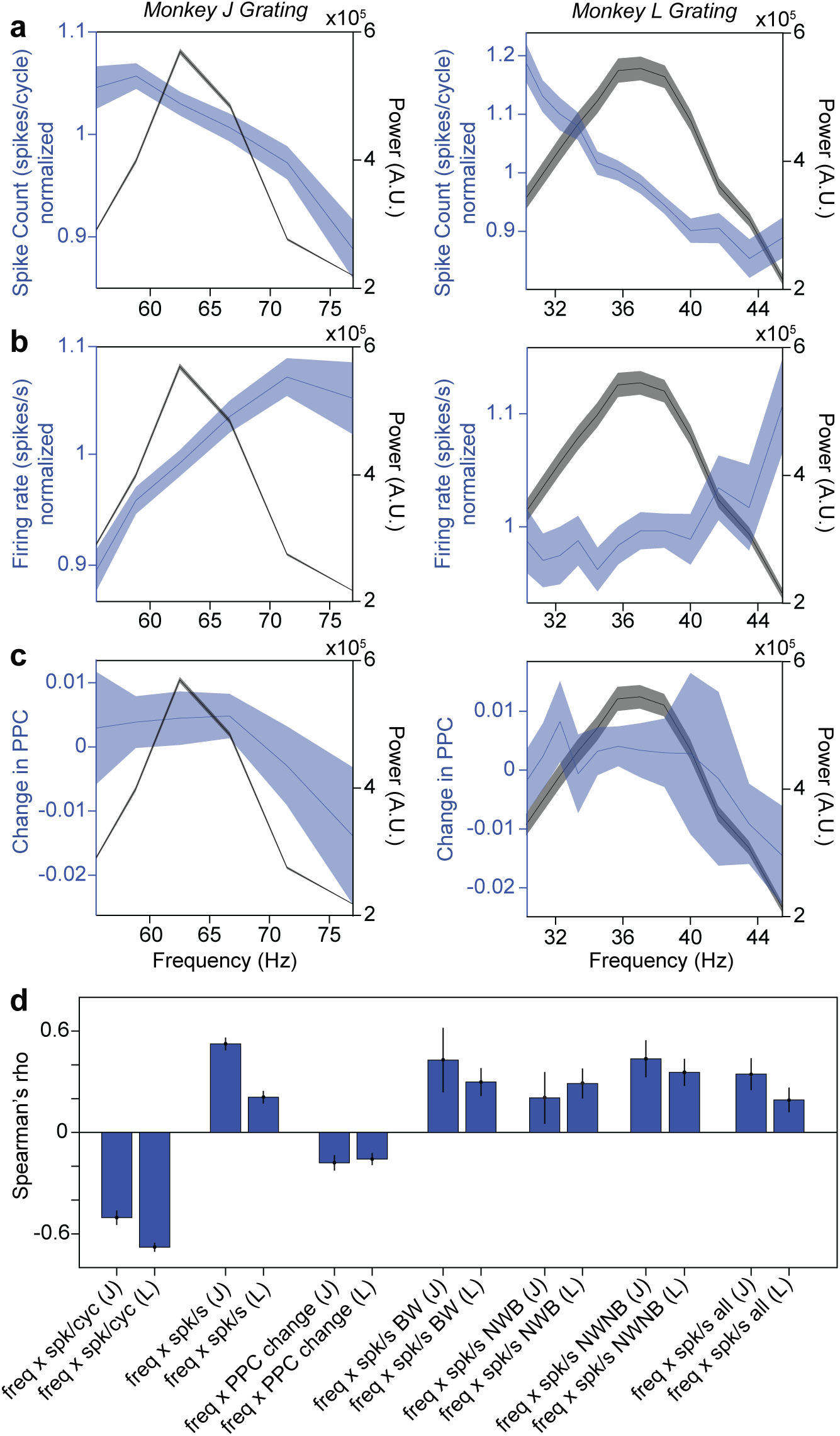
The relationship between gamma-cycle duration and spiking. (**a**) The blue curve depicts the average normalized multi-unit (MU) spike count in detected gamma cycles of different durations, expressed on the x-axis as frequencies, for monkey J (left) and monkey L (right). The black curve depicts raw power in the gamma range of the respective monkeys. Error regions show ±2 SEM across units. (**b**) Same as **a**, but using the normalized MU firing rate. (**c**) Same as **a**, but showing the normalized change in spike-LFP PPC. (**d**) Correlation between the gamma-cycle duration, expressed as frequency, and several spiking metrics, separately for the two monkeys (J and L). Vertical lines depict ±2 SEM across units.

We further wondered how spike synchrony was related to gamma-cycle duration. To investigate this, we (1) computed the duration of each gamma cycle, (2) identified all cycles of a certain duration, (3) pooled all spikes that were fired in those cycles together, and (4) computed spike-LFP phase-locking for each pool of spikes. We quantified phase locking with the pairwise phase consistency (PPC1) ^47^ metric, which removes potential biases due to spike count or firing history effects. We found that spike-LFP phase locking was negatively correlated with gamma frequency (Fig. 6c,d), i.e. positively correlated with gamma-cycle duration. The stronger spike-LFP phase locking in longer gamma cycles may have been due to a stronger spiking transient (at the “preferred” gamma phase), despite lower average firing-rates. To examine this, we divided each gamma cycle into eight non-overlapping phase-bins and computed MUA firing rates for these different bins. We did this separately for gamma cycles of different durations. As expected, longer gamma cycles showed a stronger phase modulation of firing rates (Fig. 7a,b). However, we did not observe a stronger spiking transient in longer gamma cycles. Instead, in longer cycles, there was a stronger suppression of firing at the “non-preferred” gamma phase.

**Figure 7.**
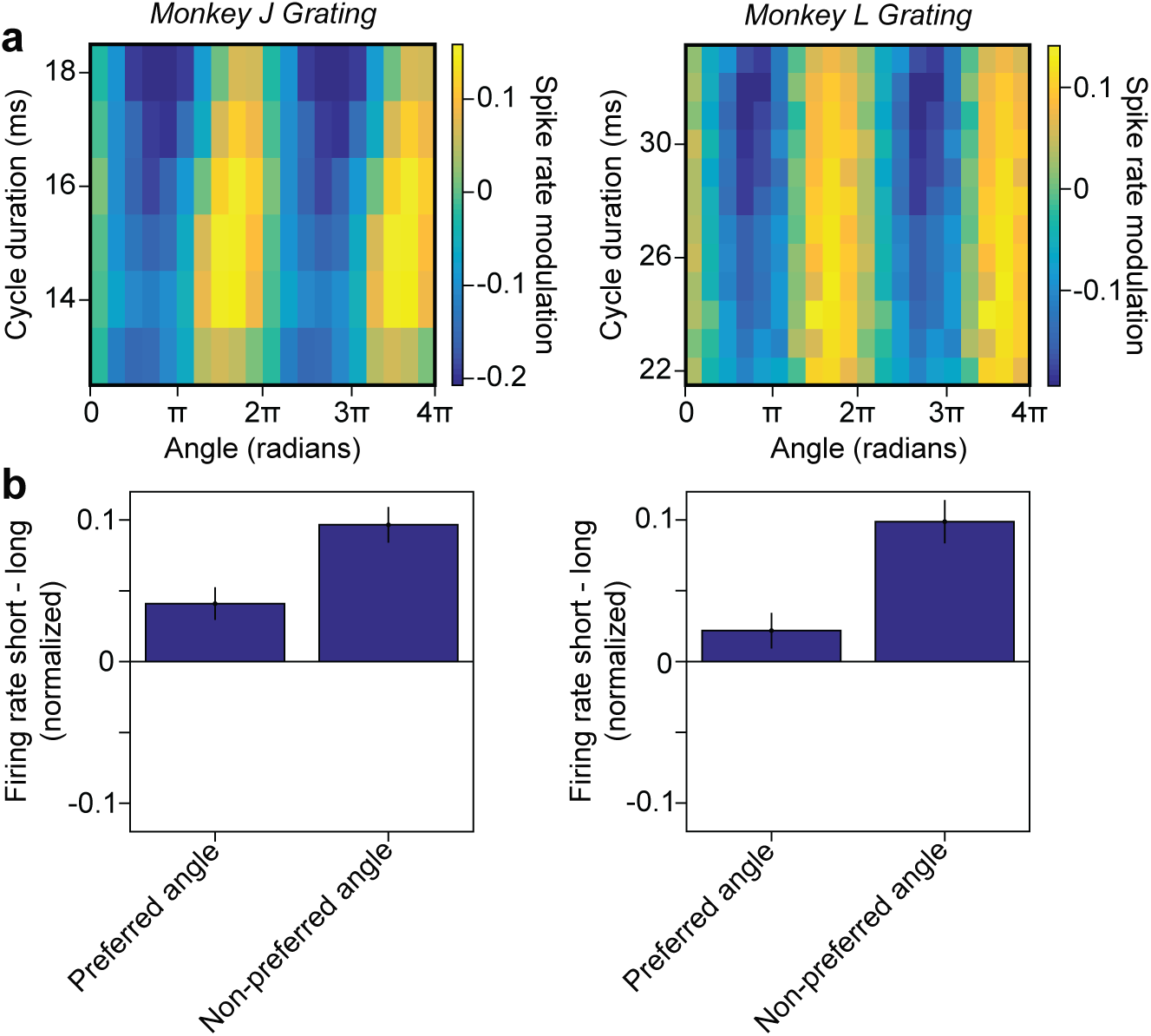
The modulation of spiking activity by the phase of the gamma cycle. (**a**) The colormap shows the modulation of the MU firing rate as a function of gamma-cycle duration (y-axis) and the phase in the gamma cycle, at which spikes occurred (x-axis). (**b**) Difference in normalized firing rate between short and long gamma cycles for the preferred (left bar) and non-preferred phase in gamma cycles (right bar). Vertical lines depict ±2 SEM across units. Data from monkey J and monkey L are shown in the left and right column, respectively.

Thus, in longer gamma cycles, average firing rates were lower, but synchrony was enhanced. This was primarily accounted for by a decrease in firing at the non-preferred gamma-phase, rather than an increase in firing at the preferred gamma-phase. These results differ from the predictions of the Atallah and Scanziani (2009) model.

### Gamma modelled by a harmonic oscillator driven by stochastic noise

Our results thus far show that correlations between gamma-cycle amplitudes and durations in awake macaque V1 are much weaker than predicted by the PING model of Atallah and Scanziani (2009). Furthermore, the relation of instantaneous firing rate with gamma-cycle duration suggests that different mechanisms are at play in primate V1. We thus wondered if a different model could explain our observations. As a starting point for developing such a model, we used our observation (Supplementary Fig. 3) that positive correlations between gamma-cycle amplitudes and durations were also present for signals generated by the stationary AR model that we fitted to the LFP data (AR contained 50 to 100 linear terms). This observation was surprising for two reasons: 1) In an AR model, all variability in amplitude and duration is due to stochastic fluctuations in the innovation term (white noise); 2) In the AR model, all the interaction terms are linear (i.e. x[t] is a linear function of past values of x[t] plus white noise), whereas previous work used models including non-linear interaction terms to produce positive amplitude-duration correlations^31^.

To generate oscillatory behavior in an AR model, the minimum number of parameters that is required is two (AR(2)). The characteristic behavior of an AR(2) can be described by its eigenvalue. When it has a complex eigenvalue, then the AR(2) model corresponds to a linear, dampened harmonic oscillator that is stochastically driven (forced) by white noise (Fig. 8a). The strength of the oscillation can be controlled by changing the magnitude of the eigenvalue (which needs to lie within the unit circle; we refer to this magnitude simply as the eigenvalue) (Fig. 8b; see Methods). We investigated whether such a simple AR(2) model produces positive correlations between gamma-cycle amplitudes and durations. To directly compare AR(2) models to the LFP gamma oscillations, we fitted AR(2) models to the LFP power spectrum, by minimizing the squared error in the gamma frequency-range. We found that AR(2) model fits could accurately reproduce the LFP power spectra in the gamma-frequency range (Fig. 8c). The eigenvalues of these fits ranged approximately between 0.97 and 0.995 (Fig. 8d); interestingly, this indicates that gamma oscillations in our V1 data were close to criticality (i.e. network instability). Next, we generated time series based on the AR(2) model and applied our method to detect gamma half-cycle amplitudes and durations. In these synthetic AR(2) signals, we observed positive correlations between amplitudes and durations (Fig. 8e). For the same range of eigenvalues, these correlations were comparable to the ones found in the V1 LFP data. Hence, positive correlations between gamma-cycle amplitudes and durations can be reproduced by a linear AR(2) model.

**Figure 8.**
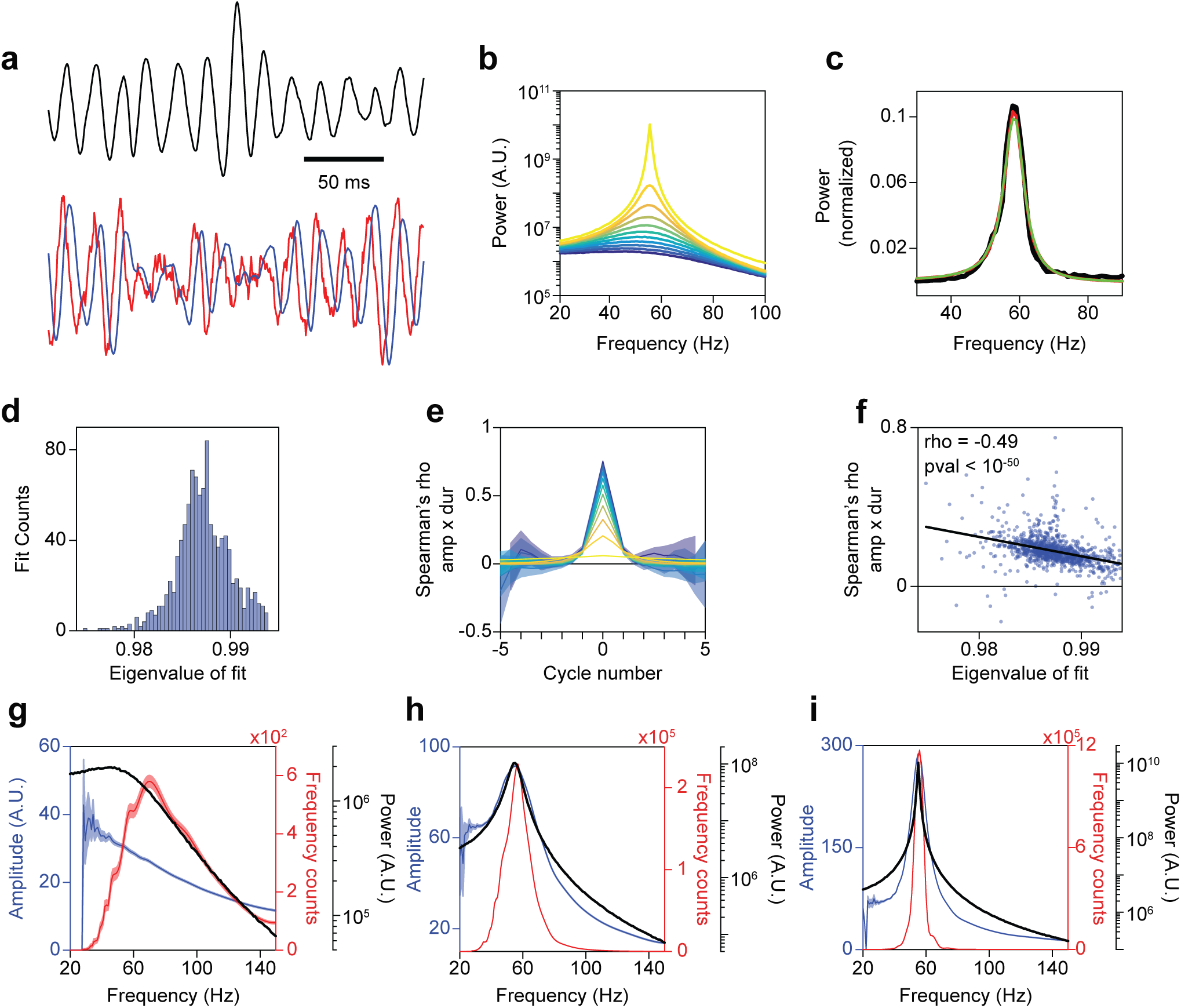
A linear harmonic oscillator driven by noise reproduces the correlation between cycle-amplitude and duration in the LFP data. (**a**; upper panel) Synthetic trace generated from a second-order autoregressive model (AR(2)). (**a**; lower panel). From the AR(2), we derived the excitatory component (red) and inhibitory component (blue) of a linear PING model with the same peak frequency and eigenvalue as the AR(2) model (see Methods). Note the characteristic delay between excitation and inhibition. (**b**) Power spectra of synthetic signals generated from AR(2) processes with corresponding eigenvalues ranging from 0.9 (dark blue) to 0.999 (bright yellow). Note that we used a periodogram with a rectangular taper, in order to minimize the spectral leakage around the peak; this can introduce an amount of broad-band leakage. (**c**) Black: The change in LFP power relative to baseline as a function of frequency (Hz), for an example site in monkey T during the presentation of a full-screen drifting grating. Red: Power spectrum of a synthetic signal generated by an AR(2) model. The AR(2) model was fitted to the LFP spectrum shown in black (see Methods). Green: Power spectrum of the I component of a linear PING model which is equivalent to the AR(2) model. (**d**) Histogram of eigenvalues corresponding to AR(2) model fits of the LFP data. (**e**) Correlation between the amplitude of a gamma half-cycle and the duration of 10 gamma half-cycles before and after it. These correlation coefficients were computed for synthetic signals generated from AR(2) processes with corresponding eigenvalues ranging from 0.9 (dark blue) to 0.999 (bright yellow) in steps of approximately 0.01. The error regions show ±2 SEM based on a bootstrap procedure. (**f**) Scatter plot of the eigenvalues displayed in **d** and the instantaneous correlation between gamma half-cycle amplitude and duration from the corresponding LFP data. The regression fit (black line) was computed with the least-squares method. (**g**-**i**) Same as **Fig. 5a**, but for synthetic signals generated from AR(2) processes with respective eigenvalues of 0.9 (**g**), 0.9871 (**h**; same as median of **d**), and 0.999 (**i**).

Based on this AR(2) model we made one further prediction, namely that correlations between gamma-cycle amplitudes and durations should be smaller when gamma oscillations are on average stronger (i.e. have a higher eigenvalue; Fig. 8e). We tested this prediction as follows: We first fitted an AR(2) model to the LFP spectra separately for each channel and condition, and determined the eigenvalue of the AR(2) fits (i.e. the oscillation strength) (see Methods). For the same LFP data, we then computed the amplitude-duration correlations, similar to Fig. 4. We then regressed the amplitude-duration correlation onto the eigenvalue of the AR(2) fits (Fig. 8f). We found that, as predicted, amplitude-duration correlations decrease as a function of the eigenvalue.

We further wondered if the CBAS of the AR(2) model would be comparable to the one of the LFP data (Fig. 5). To this end, we generated time series for AR(2) models of different eigenvalues (Fig. 8g-i). When the AR(2) eigenvalue was comparable to the one found in the LFP data (Fig. 8h), we observed a non-monotonic relationship between gamma-cycle amplitudes and gamma-cycle frequency, and a steep decline in gamma-cycle amplitudes towards higher gamma-cycle frequencies. This matched our findings for the V1 LFP data (shown in Fig. 5). Furthermore, we found that the cycle-frequency distribution was roughly symmetric around the peak gamma-frequency in the FBPS (Fig. 8g-i); similar to what we had observed for the V1 LFP data (Fig. 5). Together, these findings indicate that a simple AR(2) model predicts the observed amplitude-duration correlation and its negative dependence on average oscillation strength, as well as the joint amplitude-duration distribution.

These findings were surprising to us: We had expected that to reproduce these features from a model, a large number of parameters and variables containing non-linear interaction terms would have been required. This becomes less puzzling, however, if one considers that there exists a basic linear PING model that is mathematically equivalent to the AR(2) model (for proof see Methods; Fig. 8a). This basic PING model has the following features: It does not contain non-linear interaction terms; it only assumes stochastic input drive to the excitatory population; and it does not contain mutual inhibitory connections and mutual excitatory connections. This PING model also reproduces the characteristic time delay between the excitatory and inhibitory population as well as E/I balance (Fig. 8a).

Based on the AR(2) model, we made two more predictions concerning the variability in gamma-cycle amplitudes and durations: (1) Amplitudes should be highly correlated across gamma cycles, i.e. there should be a very high autocorrelation of the gamma-cycle amplitude. This amplitude autocorrelation should be higher when gamma oscillations are on average stronger (Fig. 9a). We determined the amplitude autocorrelation by detecting the amplitude of all detected half-cycles in the LFP data. We then computed the autocorrelation between the amplitude of a given half-cycle with the amplitude of the previous and succeeding half-cycles. We found very high autocorrelations in half-cycle amplitude that were comparable to the ones observed in the AR(2) time series (Fig. 9b,c). Moreover, we found that, as predicted, the amplitude autocorrelation was an increasing function of the eigenvalue. To rule out that the high amplitude correlations resulted from using half-cycles, we repeated this analysis on full cycles, and found essentially the same result (Supplementary Fig. 5).

**Figure 9.**
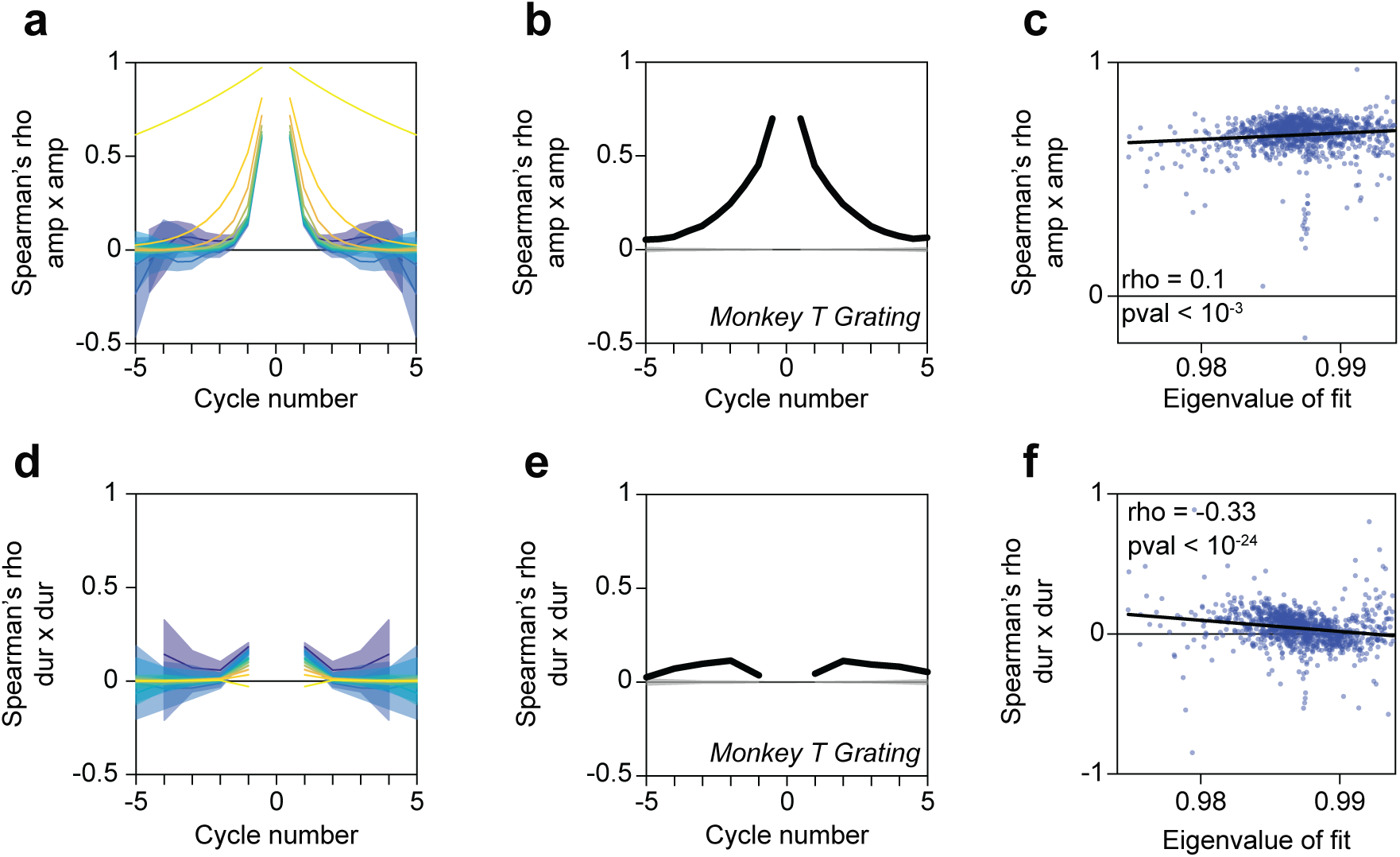
A linear harmonic oscillator reproduces gamma-cycle amplitude and duration autocorrelations in the LFP. (**a**) The correlation between the amplitude of a given gamma half-cycle and the 10 gamma half-cycles before and after it (i.e. the autocorrelation) for synthetic signals generated from AR(2) processes. The error regions show ±2 SEM based on a bootstrap procedure. (**b**) The autocorrelation for LFP data from monkey T during presentation of a full-screen drifting grating. The gray lines and gray-shaded areas depict the means and 99.9% confidence regions, after randomizing the order of duration values across trials. (**c**) Same as **Fig. 6c**, but now showing the correlation between the amplitude of a given gamma half-cycle and the amplitude of its preceding and succeeding half-cycle, pooling data points from multiple datasets, conditions and channels. (**d**-**f**) Same as **a**-**c** but for gamma full-cycle durations.

(2) The second prediction was that gamma-cycle durations should be weakly correlated across gamma cycles, especially when gamma oscillations are on average stronger (Fig. 9d and Supplementary Fig. 6a). We computed autocorrelations based on the duration of all detected half-cycles in the LFP data. We found that, as predicted, the autocorrelation of the half-cycle durations was a decreasing function of the eigenvalue (Supplementary Fig. 6c). Yet, we observed that the autocorrelations of half-cycle durations were consistently negative, different from the autocorrelations in the AR(2) time series (which were positive or close to zero) (Supplementary Fig. 6a). This feature was likely due to asymmetric wave shapes of gamma cycles, which may perhaps reflect a difference in the time constants of the AMPA and GABA currents that generate the LFP and contribute to different parts of the gamma cycle. To avoid the potential influence of cycle asymmetry, we therefore repeated our analysis for full-cycle durations. We found that, as predicted, the autocorrelations of the full-cycle duration were positive but close to zero (bootstrap mean = 0.041; bootstrap SEM = 0.0038) (Fig. 9d-f) and that the autocorrelations were a decreasing function of the oscillation strength. In essence, this means that for strong oscillations, the variability in the duration of the next gamma cycle cannot be accurately predicted from the variability in the duration of the current gamma cycle.

## DISCUSSION

Circuits of excitatory and inhibitory neurons can generate rhythmic activity in the gamma frequency-range (30-80Hz). Individual gamma-cycles show ample spontaneous variability in amplitude and duration. The mechanisms underlying this variability are not fully understood. We recorded local-field-potentials (LFPs) and spikes from awake macaque V1, and developed a noise-robust method to detect gamma-cycle amplitude and duration. We show that this method circumvents several problems that could arise due to band-pass filtering and peak/trough detection (Figure 2-3). This method allowed us to analyze the precise way in which amplitude and duration vary between gamma cycles, and how this variation relates to neuronal spiking activity. These analyses reveal several properties of gamma-oscillatory dynamics in our data:

1. The amplitude and duration of individual gamma cycles showed a weak but positive correlation (Spearman’s rho = 0.199).
2. Correlations between amplitude and duration decreased when gamma oscillations were on average stronger.
3. Gamma-cycle amplitude was strongly autocorrelated across cycles, especially for gamma oscillations that were on average stronger. Thus, if a given gamma cycle had a higher (lower) amplitude, then the next gamma-cycle also tended to have a higher (lower) amplitude.
4. Gamma-cycle duration was very weakly autocorrelated across gamma cycles, especially for gamma oscillations that were on average stronger. This implies that variability in the duration of the next gamma cycle (which would be around 10ms for a bandwidth of 40-60Hz) cannot be accurately predicted from the variability in the duration of the current gamma cycle.
5. Longer gamma cycles were associated with stronger spike-field phase-locking (synchrony), but lower firing-rates. Furthermore, longer gamma cycles are not accompanied by stronger, transient spiking activation.

We find that the first four properties can be reproduced by a linear harmonic oscillator driven by stochastic noise (AR(2) model with complex roots). We show that this model can be accurately fitted to V1 LFP data and is equivalent to a basic, linear PING (Pyramidal Interneuron Network Gamma) circuit. This basic PING model does not contain non-linear interaction terms; it only has stochastic input drive to the excitatory population; and lacks recurrent inhibitory connections as well as recurrent excitatory connections. This PING model also reproduces the characteristic time delay and balance between the excitatory and inhibitory population. We note that the idea that oscillations in the brain can be modelled as harmonic oscillators was introduced many decades ago by ^48-50^.

Our study was motivated by a previous study of Atallah and Scanziani (2009), who reported a strong positive correlation (r = 0.61) between gamma-cycle amplitude and duration in rat hippocampus. Here, we show that these positive correlations can arise due to the employed analysis method, mainly due to the presence of noisy fluctuations in the signal (Fig. 2d,g). To avoid this problem, we developed an algorithm for the detection of gamma-oscillatory epochs, i.e. periods in the LFP dominated by gamma oscillations. The correlations computed for these periods remained positive, but were substantially weaker (Spearman’s rho = 0.199; comparable result for Pearson’s r) compared to ^31^. This highlights that the detection of gamma cycle amplitude and frequency is difficult, because of the presence of non-stationarities in the analyzed signal, and filter-generated smearing between adjacent data points in the time domain. This does not mean that our method detects the “ground-truth” gamma-cycle amplitude or duration: These quantities do not describe statistical properties of the signal, in contrast to quantities like the power spectral density. In a linear harmonic oscillator driven by noise, the notion of a “cycle” becomes fuzzy for low durations and amplitudes: Fluctuations become noise-driven, and the Hilbert-transform can yield negative frequencies, i.e. phase slips. For this reason, our cycle-detection method explicitly rejects epochs with phase slips (similar to ^51^).

As we will discuss now, we reach a different conclusion about the underlying mechanisms of amplitude-duration correlations than Atallah and Scanziani (2009), although the model that we propose shares many features with their model. Before doing so, we first briefly mention several points of debate about the mechanisms of gamma oscillations. First, it is unclear in which circuits, and under which conditions, gamma oscillations can be generated, and whether they are generated by an ING (Interneuron Network Gamma) or a PING mechanism ^5, 10, 32-37^. Several studies have observed a delay between the activity of excitatory and inhibitory neurons (or intracellular E/I currents), consistent with the PING mechanism ^37, 52-56^. However, not all studies find such a phase delay ^27, 57, 58^. Moreover, both PING and ING models can produce a wide range of dynamics depending on the specific parameter settings ^36^. A second point of contention is that the relative contributions of SOM+ and PV+ interneurons remain unclear ^10, 59^. And third, in primate and cat V1, there are specialized excitatory neurons that may play a role in generating high-amplitude gamma oscillations ^38, 45^.

Here, we show that many features of gamma-oscillatory dynamics in awake macaque V1 are predicted from a surprisingly simple, stationary model containing only linear dynamics. It is often assumed that variability in gamma-cycle amplitudes and durations results from non-linear dynamics or non-stationarities in the underlying signal, e.g. due to eye movements ^26, 29^ or cross-frequency coupling ^60^. However, we show that spontaneous variability in amplitude and duration is consistent with an underlying AR(2) model that is stationary. We further show that the AR(2) model is equivalent to a linear PING model driven by stochastic inputs to the E population. This model, while sharing several features of the model by Atallah and Scanziani (2009), does not require the presence of a strong, transient bout of excitatory activity to produce long gamma cycles, as was supposed by the PING model of Atallah and Scanziani (2009). This agrees with our result that longer V1 gamma cycles are not accompanied by a stronger spiking transient (Fig. 7).

Our model connects two lines of research on gamma dynamics: On the one hand the PING model, which directly models the interaction between neuronal populations. Our linear PING model can be considered a reduced case of the linear noise approximation of the Wilson-Cowan model ^61, 62^. On the other hand, the model of gamma as filtered white noise ^7^, which, like the AR(2) model, is also a stationary signal model that reproduces the power spectrum of the signal. (Note that while the AR(2) is a form of filtered white noise, the reverse is not necessarily the case). Burns et al. showed that the distribution of gamma-burst durations can be reproduced by generating filtered white-noise, i.e. a mix of sinusoids with random phases and the same amplitude as the LFP power spectrum (which is different from an AR(p) model) ^7^. Further, by computing auto-coherence over the wavelet transform of the LFP signal, Burns et al., found relatively weak auto-coherence of gamma over time (around 0.3-0.4 resultant length) ^63^. Here we performed a similar analysis with a cycle-by-cycle detection method that avoids spurious correlations due to windowing or band-pass filtering. In our data, we find that the correlation between the full-cycle-duration of the current and the next cycle is close to zero (bootstrap mean = 0.0406), and approaches zero for strong oscillations. It remains unclear whether our very simple model reproduces all features of gamma-oscillatory dynamics; it is possible that more complex models are needed in order to do so, and our model primarily models spontaneous gamma dynamics. However, it is quite surprising that the gamma oscillations in the collective, high-dimensional dynamics of millions of V1 neurons, measured at the macro/meso-scale, are well predicted from a model that is linear and contains only two parameters.

## ACKNOWLEDGEMENTS

We thank Michael Schmid and Richard Saunders for planning and performing surgical implants, and Thomas Stieglitz and Eva-Maria Fiedler for producing the polyimide-based ECoG arrays. We thank Boris Gutkin and Gregory Dumont for helpful comments. GS was supported by ERC Starting Grant (SPATEMP to MV). MV acknowledges grant support from BMF (Computational Life Sciences) and ERC Starting Grant (SPATEMP). PF acknowledges grant support by DFG (SPP 1665 FR2557/1-1, FOR 1847 FR2557/2-1, FR2557/5-1-CORNET, FR2557/6-1-NeuroTMR), EU (HEALTH-F2-2008-200728-BrainSynch, FP7-604102-HBP, FP7-600730-Magnetrodes), a European Young Investigator Award, NIH (1U54MH091657-WU-Minn-Consortium-HCP), and LOEWE (NeFF). MLS acknowledges grant support by HFSP (HFSP fellowship LT000904/2011-L).

## AUTHOR CONTRIBUTIONS

Conceptualization, G.S., M.V., and P.F.; Methodology, G.S., M.V., and P.F.; Software, G.S., J.R.D., and M.V.; Analysis of LFP data, G.S. and M.V.; Analysis of spiking data, M.V. and I.O.; Simulations and mathematical analysis of AR(2) model, GS and M.V.; Experiments G.S., J.R.D., M.L.S., C.A.B., B.L., A.P., J.K.-L., R.R., S.N., W.S., and P.F.; Writing, G.S., M.V., and P.F.; Supervision, M.V. and P.F.; Funding Acquisition, W.S., M.V., and P.F..

## METHODS

### Subjects

We analyzed data from a total of 6 adult macaque monkeys (*macaca mulatta*), referred to as monkey H, I, J, L, P and T. Monkeys I and L are/were female, the others male. The experiments were approved by the responsible regional or local authority, which was the Regierungspräsidium Darmstadt, Germany, for monkeys H, I, J, L and T, and the ethics committee of the Radboud University, Nijmegen, Netherlands, for monkey P.

### Recordings

We used different recording procedures and stimulus paradigms for the different monkeys, and will describe these separately for the different monkeys.

### Task

All monkeys performed a passive fixation task. The specific details of the task performed by monkeys I and P were as follows: Monkeys initiated a trial by depressing a lever (monkey I) or touching a bar (monkey P), which triggered the appearance of a fixation point, and then brought their gaze into a fixation window around the fixation point. Monkeys were required to fixate on the fixation point, which was centered on a gray background, after which a stimulus was presented. If they kept their gaze within the fixation window as long as the stimulus was presented, they were given a juice reward after the release of the lever/bar following stimulus offset. Monkeys H, J, L and T performed a similar task, with the initiation/termination of the trial being solely dependent on the acquisition/release of fixation (i.e. not dependent on pressing a lever or touching a bar). Further details of this version of the task are described in ^42^ for monkey H, and in ^24^ for monkeys J and L. For all monkeys, fixation windows ranged between 0.5 and 1.2 degrees radius.

### Recordings (electrodes, reference)

For monkey H, recordings were done with CerePort (“Utah”) arrays (64 micro-electrodes; inter-electrode distance 400 μm, tip radius 3-5 μm, impedances 70-800 kΩ, half of them with a length of 1 mm and half with a length of 0.6 mm, Blackrock Microsystems). A reference wire was inserted under the dura toward parietal cortex. Further details are reported in ^42^. For monkey I, a semi-chronic microelectrode array micro-drive was implanted over area V1 of the left hemisphere (SC32-1 drive from Gray Matter Research; 32 independently movable glass insulated tungsten electrodes with an impedance range of 0.5-2 MΩ and an inter-electrode distance of 1.5 mm, electrodes from Alpha Omega). We used the micro-drive chamber as the recording reference. For monkeys J and L, recordings were performed with 2 to 10 microelectrodes, made of quartz-insulated, tungsten-platinum material (diameter: 80 μm; impedances between 0.3 and 1MΩ; wire from Thomas Recording). These were inserted independently into the cortex via transdural guide tubes (diameter: 300μm; Ehrhardt Söhne), which were assembled in a customized recording device (designed by S.N.). This device consisted of 5 precision hydraulic micro-drives mounted on an X-Y stage (MO-95, Narishige Scientific Instrument Laboratory, Japan), which was secured on the recording chamber by means of a screw mount adapter. Inter-electrode distance ranged between 1 and 3 mm. We used the micro-drive chamber as the recording reference. Further details are reported in ^24^. For monkey P, we recorded neuronal activity with a micro-machined 252-channel electrocorticogram (ECoG) electrode array implanted subdurally on the left hemisphere ^64-66^. We used a silver ball implanted over occipital cortex of the right hemisphere as the recording reference. For monkey T, we recorded neuronal activity with a micro-machined 252-channel ECoG electrode array implanted subdurally over areas V1 and V4 of the left hemisphere (252 electrodes; inter-electrode distance 1400 μm; electrode diameter 400 μm, IMTEK & BCF, University of Freiburg) ^64^. We used an electrode adjacent to the lunate sulcus as a recording reference for the section of the array covering area V1.

### Recordings (acquisition,filtering)

For monkeys H, I and T, we acquired data with Tucker Davis Technologies (TDT) systems. Data were filtered between 0.35 and 7500 Hz (3 dB filter cutoffs) and digitized at 24,414.0625 Hz (TDT PZ2 preamplifier). For monkeys J and L, we obtained spiking activity and the LFP by amplifying 1000 times and band-pass filtering (0.7-6.0 kHz for MUA; 0.7-170 Hz for LFP) with a customized 32-channel Plexon pre-amplifier connected to an HST16o25 headstage (Plexon Inc., USA). Additional 103-fold signal amplification was performed by onboard amplifiers (E-series acquisition boards, National Instruments, USA). For monkey P, we acquired data with a Neuralynx system. Data were amplified 20 times, high-pass filtered at 0.159 Hz, low-pass filtered at 8 kHz, and digitized at 32 kHz by a Neuralynx Digital Lynx system.

### Receptive field mapping/Eccentricities

Receptive fields (RFs) were mapped with either bar stimuli (^24, 42^; monkeys H, I, J, L), patches of moving gratings (^65^; monkey P) or red dots (monkey T). The signal used for RF mapping was multi-unit activity (MUA) for monkeys H, I, J, L, and the LFP gamma power for monkeys P and T. For monkeys J and L, we recorded neuronal activity from the opercular region of area V1, leading to RF-center eccentricities of 2-3 deg, and occasionally from the superior bank of the calcarine sulcus, leading to RF-center eccentricities of 10-13 deg. For monkey H, RF-center eccentricities ranged between 5.2 and 7.1 deg (median RF-center eccentricity 6.2 deg). For monkey I, RF-center eccentricities ranged between 2.6 and deg (median RF-center eccentricity 4.5 deg). For monkey P, RF-center eccentricities ranged between 3 and 5.7 deg (median RF-center eccentricity 4.6 deg). For monkey T, RF-center eccentricities ranged between 3.1 and 7.1 deg (median RF-center eccentricity 3.8 deg).

### Eye position monitoring

For monkeys H, I and T, eye movements and pupil size were recorded at 1000 Hz using an Eyelink 1000 system (SR Research Ltd.) with infrared illumination. For monkeys J and L, we monitored the eye position with a scleral search coil system (DNI, Crist Instruments, USA; sampling rate of 500 Hz). For monkey P we monitored eye position with an infrared camera system (Thomas Recording ET-49B system) at a sampling rate of 230 Hz. We used a standardized fixation task in order to calibrate eye signals before each recording session.

### Behavioral control and stimulus presentation

Stimulus presentation and behavioral control was implemented as follows: The software toolbox ARCADE ((Dowdall et al., 2018) https://gitlab.com/esi-neuroscience/arcade) was used for monkeys H, I and T; Custom LabVIEW code (Lab-VIEW, National Instruments, USA) was used for monkeys J and L; The software toolbox CORTEX (dally.nimh.nih.gov/index.html) was used for monkey P.

Monkeys H and I were presented with full-screen uniform color surfaces. Surface color varied across trials according to a pseudo-random sequence. For our analyses, we used the hue that elicited the strongest gamma oscillations (monkey H RGB: 149 99 0; monkey I RGB: 255 0 0). In a separate session, monkey I was also repeatedly presented with a full-screen drifting square-wave red-and-green grating of a fixed initial phase and drift-direction (RGB for red 255 0 0 and green 0 255 0; spatial frequency: 1.5 cycles/degree; temporal frequency 2 Hz). Monkeys J and L were presented with large drifting square-wave black-and-white gratings (spatial frequencies: 1.25-2 cycles/degree; temporal frequencies: 1.4-2Hz) and plaid stimuli. Only the gratings were used for our analyses. The gratings had a diameter of 8 degrees of visual angle and were positioned at the average of the RF centers of the recorded MUA. In each trial, the direction of the grating drift was randomly chosen from 16 directions (in steps of 22.5 degrees). Monkey P was repeatedly presented with a full-screen drifting square-wave black-and-white grating of a fixed initial phase and drift-direction (spatial frequency: ∼1 cycle/degree; temporal frequency ∼1Hz). Monkey T was presented with full-screen uniform color surfaces, with the color changing across trials according to a pseudo-random sequence. For our analyses, we used two hues that elicited the strongest gamma oscillations (RGB: 255 0 0 and 0 0 255). In separate sessions, monkey T was also presented with full-screen drifting square-wave colored gratings of pseudo-random initial phases and drift-directions. For our analyses, we used the gratings that elicited the strongest gamma oscillations (red-green RGB: 255 0 0 and 0 255 0 and blue-yellow RGB: 0 0 255 and 255 255 0; spatial frequency: 1.5 cycles/degree; temporal frequency 2 Hz). For monkeys H, I and T, stimuli were presented on 120 Hz LCD monitors ^67^, without gamma correction. For monkeys J, L and P, stimuli were presented on CRT monitors (100-120 Hz), after gamma correction.

### Data analysis

All analyses were done in MATLAB (The MathWorks) using custom scripts and the FieldTrip toolbox (www.fieldtriptoolbox.org ^68^). The analyses were done only on correct trials. In monkeys P and T, we selected the 25% electrodes/sites over area V1 with the strongest visually induced gamma band activity, because the grids covered a relatively large region of retinotopic space and contained electrodes that were poorly driven by the visual stimulus. In monkeys H, I, J and L, we analyzed all visually driven electrodes. In all monkeys except for monkey T, we analyzed LFP signals that were recorded relative to the common reference signal (described above). For monkey T, we calculated local bipolar derivatives between LFPs from immediately neighboring electrodes. i.e., differences (sample-by-sample in the time domain), similar to previous studies ^65^. This was done because the global references in monkey T were positioned over V1 and V4 in the same hemisphere.

### Preprocessing

For monkeys H, I and T, LFPs were obtained from the broadband signal after low-pass filtering (sixth order Butterworth filter with a corner frequency of 500 Hz), high-pass filtering (third order Butterworth filter with a corner frequency of 2 Hz for monkey T and 4 Hz for monkeys H and I) and down-sampling to 2034.51 Hz. For monkeys J and L, LFPs were filtered between 0.7-170Hz (hardware-filter, described above) and down-sampled to 1 kHz. For monkey P, we obtained LFP signals by low-pass filtering at 200 Hz and down-sampling to 1 kHz. In addition, for monkey P, we removed powerline artifacts at 50 Hz and its harmonics with a digital notch filter.

### Segmenting Data into Epochs, and Calculation of Power and TFR

To estimate the LFP power spectra in the stimulus and baseline periods (Figs 1b,c,g,h, 5, 6a-c, and Supplementary Figs 3b, 4), we used the following procedure: Power spectra were estimated separately for the pre-stimulus period and the stimulation period. The pre-stimulus period was the time between fixation onset and stimulus onset. During the pre-stimulus period, monkeys fixated on a central dot on a gray screen, and there was no other stimulus presented. For monkeys H, I, P and T, the pre-stimulus and stimulation periods were of variable length across trials. We kept data corresponding to the pre-stimulus and stimulation period with the minimum length (monkey H: baseline 0.3s / stimulation 1.5s; monkey I: baseline 0.5s / stimulation 2s; monkey P: baseline 0.3s / stimulation 2.3s; monkey T: baseline 1.1s / stimulation with full-screen gratings 2.8s / stimulation with full-screen uniform color surfaces 3.2s). For monkeys J and L, the pre-stimulus and grating-stimulation periods had a stable duration across trials within a session but their duration varied between sessions. All of the available pre-stimulus and grating data were analyzed for those monkeys (baseline 0.8-1s / stimulation 2-2.4s). The power spectral analysis was based on epochs of fixed lengths. Therefore, the described task periods were cut into non-overlapping epochs. We aimed at excluding data soon after stimulus onset (“event”) to minimize the influence of the stimulus-onset related event-related potential on our analyses. Therefore, periods were cut into non-overlapping epochs, starting from the end of the period and stopping before an epoch would have included data approximately 0.5 s after those events. For Fig. 1b,c,g,h, the estimation of power spectra was based on epochs of 0.5 s length; for Figs 5, 6a-c and Supplementary Figs 3b and 5, power spectra were based on epochs of 0.25 s. Data epochs were Hann tapered, to achieve a fundamental spectral resolution (Rayleigh frequency) of 2 Hz (4 Hz for Figs 5, 6a-c and Supplementary Figs 3b and 5), and then Fourier transformed. The gamma-band power spectra used for the AR(2) fits (Figs 8c,d,f, 9c,f, and Supplementary Figs 5b, 6c), the power spectra of synthetic AR(2) signals (Fig. 8b), and the joint distribution of gamma-cycle amplitude and duration (Fig. 8g-i) were based on rectangular windows of 1s, in order to ensure minimal spectral smearing, and thus a more accurate fit. For the time-frequency analysis of power, we used window lengths of ±2.5 cycles per frequency which were slid over the available data in steps of 1 ms. Power during the stimulation period was normalized to the pre-stimulus baseline period, separately for each channel, in the following manner: Power per frequency and per trial was calculated as described above. Power calculated for the pre-stimulus baseline period was then averaged across trials. Finally, trial-wise normalized power was calculated for the stimulation period by subtracting the average pre-stimulus spectrum and then dividing by it.

### Spike sorting

Single units were isolated through semi-automated spike sorting ^38^. First, we performed semi-automatic clustering with the KlustaKwik 3.0 software. The energy of the spike waveform and the energy of its first derivative were used as features in this procedure. A candidate single unit was accepted if the corresponding cluster was clearly separable from the noise clusters, and if the inter-spike-interval distribution had a clear refractory-period. This was done manually with the M-Clust software. In addition, we used the isolation distance (ID; ^69^) as a measure of cluster separation. The ID of a candidate single unit had to exceed 20 in order for it to be included in our analyses. The median ID was 25.05. This procedure led to the isolation of 100 single units. For each isolated single unit, we computed the peak-to-trough duration of the average AP waveform. Single units with long (>0.235ms) and short (<0.235ms) peak-to-trough durations were named “broad-waveform” (BW) and “narrow-waveform” (NW) neurons, respectively. Broad-waveform neurons corresponded to 29% of the single unit population.

### Initial estimation of gamma-cycle amplitude and duration (cf. Atallah & Scanziani, 2009)

For our initial analyses of individual gamma cycles, we implemented the algorithm as described by Atallah and Scanziani (2009) for data from awake freely-moving rats. In short, we first low-pass filtered the LFP by using a 40 ms moving average filter and then subtracted this filtered signal from the original time series (Experimental Procedures and Supplemental Experimental Procedures of Atallah and Scanziani, and their personal communication with us), which effectively corresponds to a high-pass filter with a corner frequency at approximately 20 Hz. The resulting signal was further band-pass filtered in the range of 5-100 Hz with a 3^rd^ order, two-way Butterworth filter. Gamma-cycle peaks and troughs were then defined as local maxima and minima, respectively. Furthermore, gamma-cycle amplitudes were defined as the difference between the voltage of a given peak and its subsequent trough. Similarly, gamma-cycle durations were defined as the interval between a given peak and it subsequent peak. This analysis was done in segments of the filtered signal which displayed high power in the individual gamma frequency range of each dataset (peak gamma frequency±20 Hz). These segments were extracted in the following way: A time-power representation of each trial was calculated with 5 discrete prolate slepian sequences and windows of 100 ms which were slid over the available data in steps of 25 ms. Gamma episodes were defined as segments of the resulting time-series which lasted for more than 100 ms and had power that exceeded a threshold. This threshold was calculated separately for each trial as the difference between the mean of the time-power representation and its standard deviation.

### Generation of colored noise

In Figure 2G, we analyzed the correlations obtained with the Atallah-Scanziani method for colored noise. We generated noise with power spectra following a 1/f^n^ function, where f denotes frequency and n assumes 11 equally spaced values between, and including, 0 (corresponding to white noise) and 2 (corresponding to Brownian noise). This was done in the following manner: (i) 1000 white noise traces containing 10^6^ samples were generated for each n. (ii) Each trace was Fourier transformed. (iii) The complex coefficients of the positive frequencies in the resulting spectra were multiplied by the 1/f^n^ function. (iv) A synthetic spectrum was constructed by concatenating the above complex coefficients with the conjugate of their flipped version. (v) The resulting spectrum was inverse Fourier transformed to obtain time series.

### Improved estimation of gamma-cycle amplitude and duration

We developed an improved method to extract gamma-cycle amplitude and frequency from the LFP signals as follows:

1. We computed the Hilbert-transform of the broadband LFP signal to obtain the analytic signal and derive the time-resolved phase from it. We used the broadband signal, because band-pass filtering creates dependencies between voltage values across time points, and can transform transient, non-oscillatory deflections into rhythmic events.
2. We detected gamma cycles as follows: First, we detected all the zero-crossings of the phase. Such phase zero crossings occur in the neighborhood of peaks and troughs in the original LFP signal. For each k-th zero-crossing, we examined whether the angular velocity of the phase was positive for all time points between the k -1-th to the k + 1-th zero-crossing (similar to ^70^). If this was not the case, then there was a negative “phase-slip” in which the instantaneous frequency became negative, and the respective zero crossing plus/minus two neighboring zero crossings were discarded. Negative instantaneous frequencies make the interpretation of the instantaneous frequency and amplitude ambiguous, and are typically accompanied by small peaks/troughs in the LFP signal. This violates our model of the gamma oscillation as a signal with a positive frequency which fluctuates over time, y(t) = A(t) * cos (ω(t)*t + φ), where A(t) and ω(t) are the instantaneous amplitude and frequency fluctuating over time.

If there was no negative phase-slip, then we identified gamma peaks by first detecting negative-to-positive zero crossings in the phase of the analytic signal. For each of these crossings, we then identified the nearest local maximum in the LFP signal (Fig. 3d). Likewise, gamma troughs were identified by detecting positive-to-negative zero crossings and identifying nearby local minima. Using the detected gamma peaks and troughs, we then determined the gamma-cycle amplitude and duration. To obtain estimates of gamma-cycle amplitude and duration with the maximum attainable temporal resolution, we divided each gamma cycle into “half-cycles”: The first half-cycle comprised the data segment from the trough to the peak, and the second half-cycle from the peak to the trough. For each half-cycle, amplitude was defined as the difference between the respective peak and trough, and duration was defined as the corresponding time interval. For each detected half-cycle, we thus obtained an amplitude and duration value. For comparison, we also determined amplitude and duration for full gamma cycles. A gamma cycle comprised the data from one peak to the next peak. Amplitude was defined as the voltage difference between the first peak and the trough. Duration was defined at the time between the two peaks.

Note that for the analysis of the relationship between individual gamma cycles and spiking activity, we used a band-pass filter (3^rd^ order, two-pass Butterworth, with a pass-band of 40-90 Hz for monkey J and 25-55 Hz for monkey L). In this case, we used an additional criterion to reject epochs of spurious oscillatory activity ^38^: We ran the same cycle-selection procedure on the pre-stimulus period, in which narrow-band gamma-band oscillations are virtually absent. For the pre-stimulus period, we obtained the mean μ_pre_ and standard deviation σ_pre_ of the distribution of amplitudes. These amplitudes were measured as the peak-to-trough distance of the gamma cycle. A cycle in the stimulus period with amplitude A was only selected if (A – μ_pre_)/σ_pre_ > 1:63 (which is equivalent to a one-sided T-test at P < 0.05). We filtered the LFP with the purpose of increasing the number of selected gamma epochs, considering that the analysis of unit firing rates and spike-field phase-locking demands a relatively large amount of data. Note that we have shown in Fig. 5 that the distributions of amplitude and frequency after band-pass filtering are comparable to the distributions obtained without band-pass filtering. In addition, the potential issues related to filtering only apply to the calculation of correlations of amplitude and duration and not to the calculation of the correlation of spiking strength and gamma frequency. This is due to the fact that filtering may generate artificial correlations between the amplitudes and durations of deflections of the same time series (explained further in the results section). The filter used on the LFP is not used on the spiking activity. Thus, artificial correlations between spiking and cycle-by-cycle frequency are not likely.

Amplitude and frequency values were extracted from selected gamma epochs of a duration of at least 2 full cycles.

### Computation of time-resolved correlations between amplitude and frequency

In the case of our V1 recordings, we observed that gamma amplitude and cycle duration progressively increased over time after the onset of a drifting grating stimulus. (Fig. 1c,d). By contrast, after the onset of a uniform color surface, gamma amplitude and duration progressively decreased and increased over time, respectively (Fig. 1g,h). These changes with time after stimulus onset could contribute to the correlation values between gamma-cycle amplitude and duration, if gamma amplitude and duration values are concatenated across all trials and time points. This would conceal the relationship between gamma-cycle amplitude and duration due to intrinsic variability, by introducing a positive or negative correlation bias for drifting gratings and uniform color surfaces, respectively.

We avoided these effects by using the following method: We calculated correlations between gamma-cycle amplitudes and durations across all trials, separately for each time point (at the respective sampling rate) after stimulus onset, and subsequently averaged those correlation values over time points and subsequently over recording sites. To enable this, we needed to define gamma-cycle amplitudes and durations for each time point. Therefore, each time point (relative to stimulus onset) was localized to the gamma half cycle (or full cycle), into which it fell, and it was assigned the respective amplitude and duration of that half cycle (or full cycle). For the calculation of correlations with one or multiple half-cycle (or full-cycle) lags, correlations were calculated between amplitudes and durations shifted relative to each other by the corresponding number of half-cycles (or full cycles).

In datasets containing more than one stimulus condition, correlation coefficients were calculated separately for each condition and then averaged across conditions.

As mentioned in the results section, the correlation analysis used the Spearman correlation coefficient. Like in ^31^, we found results to be essentially identical for Spearman and Pearson correlation, when using their method of determining gamma-cycle amplitude and duration. For the rest of our analyses, we used exclusively the Spearman correlation coefficient.

### Statistical significance of correlations

The statistical significance of auto- and cross-correlations of gamma-cycle amplitudes and durations, and correlations between AR(2)-fit eigenvalues and auto- or cross correlations of gamma-cycle amplitudes and durations was assessed by means of a non-parametric randomization approach. In this paragraph, we will describe this approach for the cross-correlation of amplitudes and durations: The order of valid duration values was randomly shuffled across trials, separately for each time-point. We then calculated surrogate Spearman’s correlation coefficients 1000 times as described above for each dataset. Next, we performed a fit of a Gaussian distribution on the 1000 surrogate correlation coefficients. Empirical correlations were deemed significant if they were 3 standard deviations larger or smaller than the mean of the surrogate distribution. This procedure implements a non-parametric version of a two-sided test with a p-value of ≈0.001.

To test if the mean correlation of gamma-cycle amplitudes and durations is significantly different from zero across datasets, we applied a Student’s t-test. In general, we prefer non-parametric randomization tests over parametric tests (like the t-test). However, some analyses contained only four or five datasets, which effectively precludes the application of non-parametric tests. Where possible, we supplemented the t-test with a non-parametric statistical test (Figs 2c, 4a,c, and Supplementary Fig. 1a). Specifically, we calculated the mean correlation across datasets for each possible combination of values that results after independently inverting or maintaining the sign of each correlation value (i.e. a full permutation). This led to a surrogate distribution of mean values to which the empirical mean was compared for statistical significance. Mean correlations were deemed significant if they were larger (smaller) than the top (bottom) 2.5 percentile of this surrogate distribution.

### Regression analysis

We performed regression analyses separately for gamma-cycle amplitudes and durations with the Matlab function *regress*. As explained in the results section, for each half-cycle, we regressed the amplitude value of the ongoing half-cycle against the amplitude values of the previous and next half-cycle, by using a least squares approach. We used the same procedure for half-cycle duration values. This was done for each point after stimulus onset separately, and by using all the amplitude and duration values across trials (for that time point). We then calculated the regression residuals by subtracting each amplitude and duration regression vector from the corresponding amplitude and duration values, separately for each timepoint. These residual values measured the extent to which the amplitude or duration in the ongoing half-cycle was greater or smaller than in the surrounding half-cycles, and thereby departed from slower trends. We then computed the correlation between the regression residuals for amplitude and duration, in the same way as described above.

### Micro-saccade detection

We low-pass filtered vertical and horizontal eye position signals by replacing each value with the average over itself ±15 ms. We then computed the first temporal derivative of the signals to obtain the vertical and horizontal velocities. We combined those values to obtain the eye speed irrespective of the direction of eye movement. Per trial, we determined the SD of eye speed, and any deviation >4 SDs and lasting for at least 30 ms was deemed a saccadic eye movement. Saccadic eye movements that remained within the fixation window were considered to be MSs.

### AR

In Supplementary Fig. 3, we computed our correlations for data generated through auto-regressive models with a power spectrum similar to the recorded LFP data. An autoregressive (AR) model of order n represents each value in a time-varying process as the linear sum of its n preceding values (each weighted by a separate coefficient) and a stochastic term. This model can then be used to generate a synthetic time series that has the same power spectrum as the original process, but that is devoid of higher-order statistical properties such as slow temporal trends or spectral cross-frequency dependencies. We modelled the LFP as an AR process of a relatively high order (50 for monkeys J and P, whose analysis was based on a sampling rate of 1000 Hz, and 100 for monkeys H, I, T, whose analysis was based on a sampling rate of 2034.51 Hz). We did this by fitting a vector of AR coefficients and a noise variance term with the Matlab function *arfit*, simultaneously to all the trials of a given stimulus condition and independently for each recording site. For our analyses, we only used the period of the trial starting at 250 ms after stimulus onset, thereby omitting stimulus onset-related transient activity. These AR models were then used to generate surrogate time series.

### AR(2) Model Derivation

Let *x*_*t*_ be a stationary stochastic signal (which could represent an LFP signal, for example). The AR(2) model is defined by the second order difference equation:

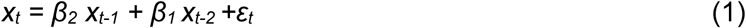

with an expected value *EV{ε*_*t*_, *ε*_*t*+*k*_*} = 0* for all time delays *k* (i.e. ε_t_ is uncorrelated white noise). For a certain range of parameters, this model is a linear, dampened harmonic oscillator driven by stochastic noise. We now rewrite this second-order difference equation into the two corresponding first-order, linear differential equations. We first swap variables and define *I = x*_*t*_ and *E = x*_*t*_ *-x*_*t-1*_. We then obtain the system of equations

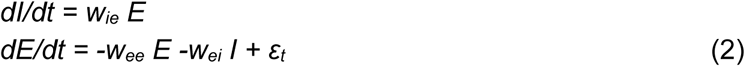

Here *w*_*ei*_ *= 1- β*_*2*_*- β*_*1*_ is the inhibitory feedback from *I* to *E*, and w_ee_ = 1 + β_1_ and w_ie_ = 1. The product *-w*_*ee*_ *E* controls the return to the steady state, in the absence of stochastic noise input. Note that the model does not contain recurrent inhibitory or excitatory connections, and does not contain any non-linear interactions. It differs from the Wilson-Cowan or PING model because of the absence of non-linearities. But it is directly related to the linear noise approximation of the stochastic Wilson-Cowan model, with the difference that there are no recurrent excitatory or inhibitory connections in the AR(2) model. From the AR(2) model, we can obtain eigenvalues in the standard way, i.e. from the roots of the characteristic polynomial equation.

To generate AR(2) signals, we computed the AR(2) coefficients for a given eigenvalue magnitude (simply referred to as eigenvalue) and oscillation frequency, using standard analytical transformations. Generated time series were analyzed with the same cycle-detection method as the LFP data. The only difference was that for the AR(2), we did not divide the data into trials, and thus computed the correlation between cycle amplitude and duration across all the cycles over all the time points (i.e. not across trials for each time point separately). In order to compare the AR models to the LFP data, we ensured that the model used a sampling frequency of 2035 Hz, similar to the sampling frequency of most of our LFP datasets. For several analyses, we correlated the eigenvalue of the AR(2) fit to the LFP data, with several correlation measures across LFP datasets, including the amplitude-duration correlation, amplitude autocorrelation and duration autocorrelation. To ensure that all preprocessing (sampling rate; filtering) was similar for these data, we only included datasets with a similar sampling frequency of 2034.51 Hz.

### AR(2) Model Fit to LFP data

We estimated the strength of gamma oscillations in our LFP data as follows: (1) We computed gamma-band power spectra separately for each channel and condition. The power spectra were based on rectangular windows of 1s, in order to ensure minimal spectral smoothing, and thus a more accurate fit. (2) We then estimated the coefficients of equivalent AR(2) models by minimizing the squared error in the gamma frequency-range (*matlab* function *fminsearch*) between each LFP power spectrum and the following function:

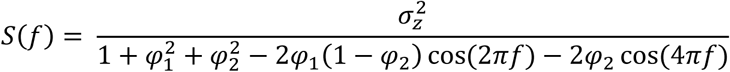

where *S(f)* is the power spectrum of the AR(2), *σ*_*z*_ is the standard deviation of this power spectrum, *f* are frequencies in the gamma range, and *φ*_1_/*φ*_2_ are the AR(2) coefficients (Fig. 8c). (3) We determined the eigenvalues of the equivalent AR(2) models (Figs 8d,f, 9c,f, Supplementary Figs 5c, 6c).

### PPC

For the calculation of spike-LFP PPC, the gamma phase of each spike within a gamma cycle was defined as t/T*2*π, where t was the time of the spike relative to the start of the gamma cycle, and T was the duration of the gamma cycle. This constitutes a linear phase interpolation. This used the improved Hilbert-based definition of gamma half-cycles (cycles). The obtained spike phases from separate trials were collected, and the average consistency of phases across these pairs was estimated with the pairwise-phase-consistency metric (PPC) ^47, 71^, and more specifically its PPC1 variant ^71^. Any potential bias due to differences in discharge rates is removed by the pairwise computation. Only neurons that fired at least 50 spikes were considered, because phase-locking estimates can have a high variance in cases of low spike counts. We were not able to perform this analysis for single-unit activity, due to the lack of a sufficient number of detected single unit spikes.

### Computation of the Cycle-Based Amplitude Spectrum (CBAS) and cycle-frequency distribution

For Fig. 5 and Supplementary Fig. 4 we computed the cycle-based amplitude spectrum (CBAS) and the cycle-frequency distribution as follows. Gamma half-cycle amplitude and duration values were extracted from the LFP through the use of the previously described improved detection algorithm. Values of gamma-half-cycle durations were converted into values of gamma-half-cycle frequency (frequency being the inverse of duration). This was done separately for each recording site and stimulus condition. Next, gamma half cycles were assigned to their corresponding frequency bin, and for each frequency bin, the average amplitude and the rate of incidence of that frequency were determined.

Note that the peak gamma-frequency varies across experimental subjects and stimulus conditions. In order to compute averages across stimulus conditions and monkeys, it is therefore necessary to align individual distributions to the power-spectral peak in the gamma-frequency-range, separately for each stimulus condition and dataset. We performed this alignment in the following way: The raw trial-wise power spectra were estimated separately for each stimulus condition as described above (see power), and from these spectra we determined the peak gamma-frequency. In addition, this was done for the baseline-corrected power spectra. The alignment of half-cycle amplitudes and frequency counts was then performed around the resulting frequency. Specifically, half-cycle amplitude and frequency count averages at ±20 Hz around the gamma peak were averaged across stimulus conditions and datasets. Note that we analyzed datasets with different sampling rates. This entailed that the range of detectable half-cycle frequencies (i.e. sampling rate/(2*duration)) varied across different datasets and, depending on the sampling rate, certain frequency bins were necessarily empty. In order to average across datasets with different sampling rates, we therefore performed a linear interpolation between normalized half-cycle amplitude values and frequency counts, which were adjacent to empty bins.

---

**Supplementary Figure 1.**
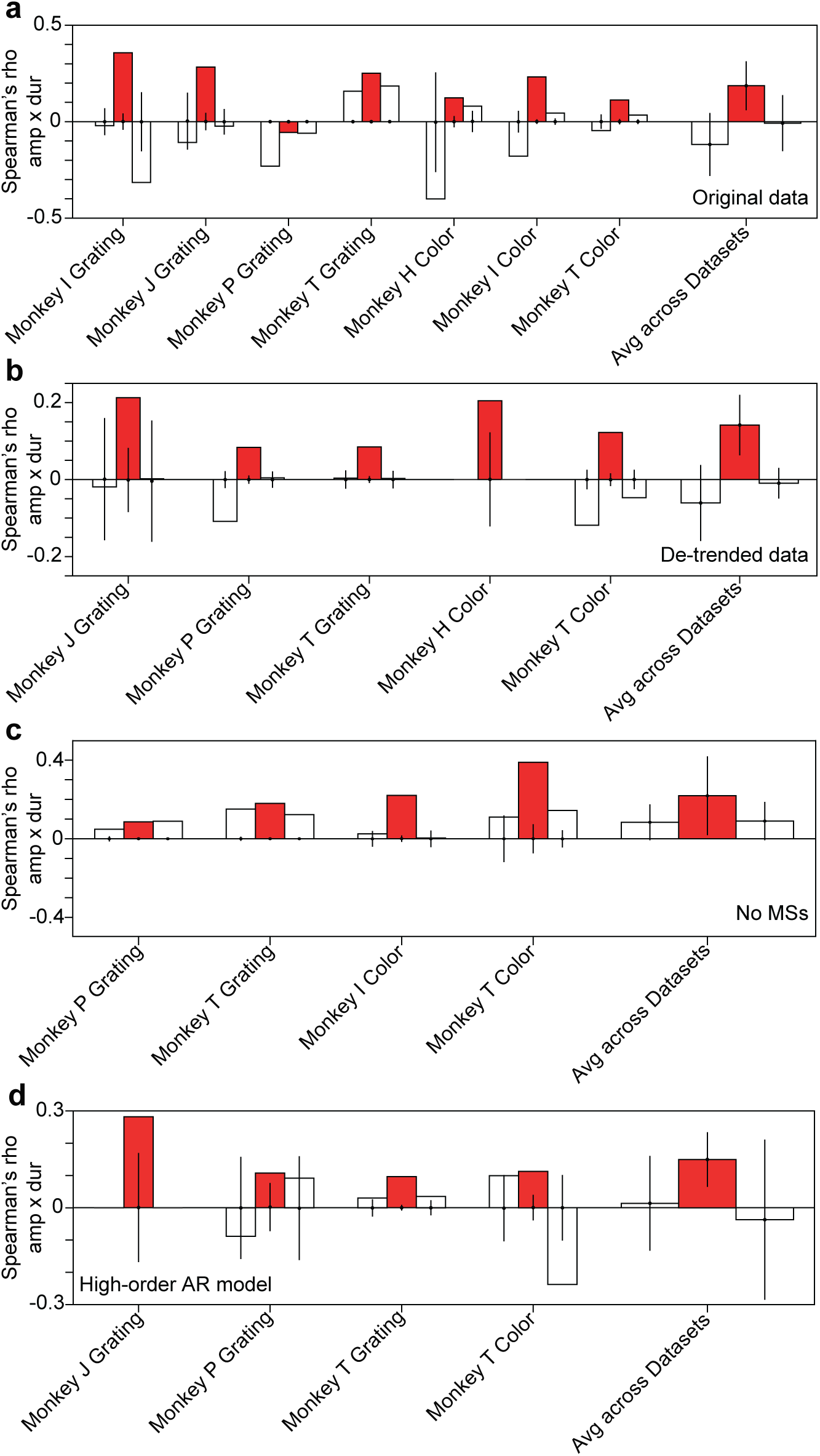
Gamma-full-cycle amplitudes and durations are positively correlated. (**a**) Same as **Fig. 4a**, but using full gamma cycles (averages across datasets: red bar: p=0.011, t-test across datasets; p<0.05, two-sided randomization test across datasets; white bars p=0.13 and p=0.9, respectively for preceding and succeeding cycles, t-test across datasets; p>0.05, two-sided randomization test across datasets). (**b**) Same as **Fig. 4c**, but using full gamma cycles (averages across datasets: red bar: p=0.008, t-test across datasets; white bars p=0.15 and p=0.51, respectively for preceding and succeeding cycles, t-test across datasets). (**c**) Same as **Supplementary Figure 2c**, but using full gamma cycles (averages across datasets: red bar: p=0.046, t-test across datasets; white bars p=0.11 and p=0.13, respectively for preceding and succeeding cycles, t-test across datasets). (**d**) Same as **Supplementary Figure 3e**, but using full gamma cycles (averages across datasets: red bar: p=0.041, t-test across datasets; white bars p=0.9 and p=0.7, respectively for preceding and succeeding cycles, t-test across datasets).

**Supplementary Figure 2.**
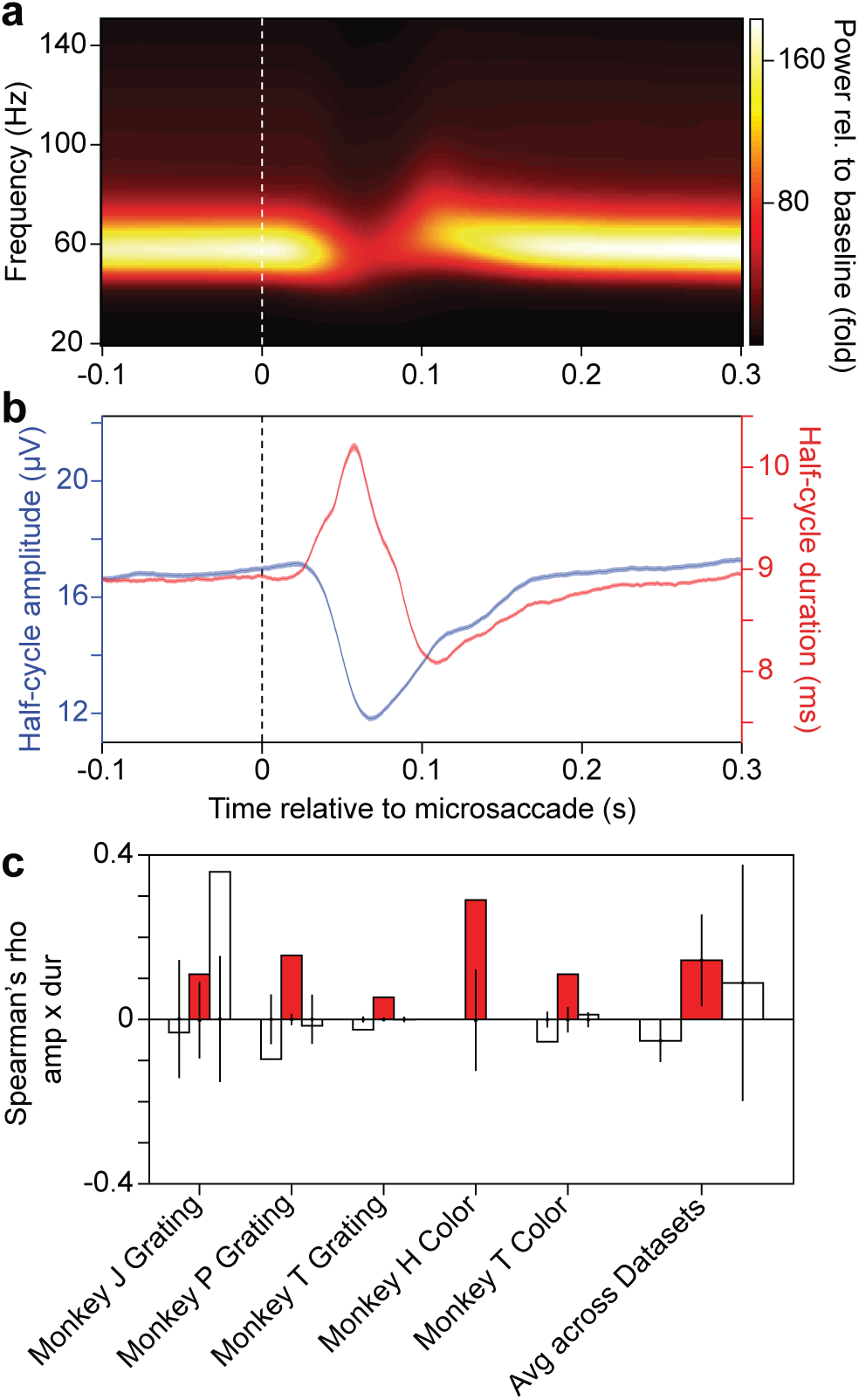
The effect of microsaccades on the correlation between gamma-half-cycle amplitudes and durations. (**a**) Time-frequency power averaged over all selected V1 recording sites in monkey T during the presentation of a full-screen drifting grating, normalized by the pre-stimulus baseline. X-axis shows time relative to detected microsaccades (MSs). (**b**) Time-course of the gamma-half-cycle amplitude (blue) and duration (red) of the data depicted in **a**. Error regions show ±2 SEM based on a bootstrap over MSs. (**c**) Same as **Fig. 4c**, but after the removal of 250 ms epochs following the occurrence of MSs for all available datasets (averages across datasets: red bar: p=0.02, t-test across datasets; white bars p=0.07 and p=0.97, respectively for preceding and succeeding cycles, t-test across datasets).

**Supplementary Figure 3.**
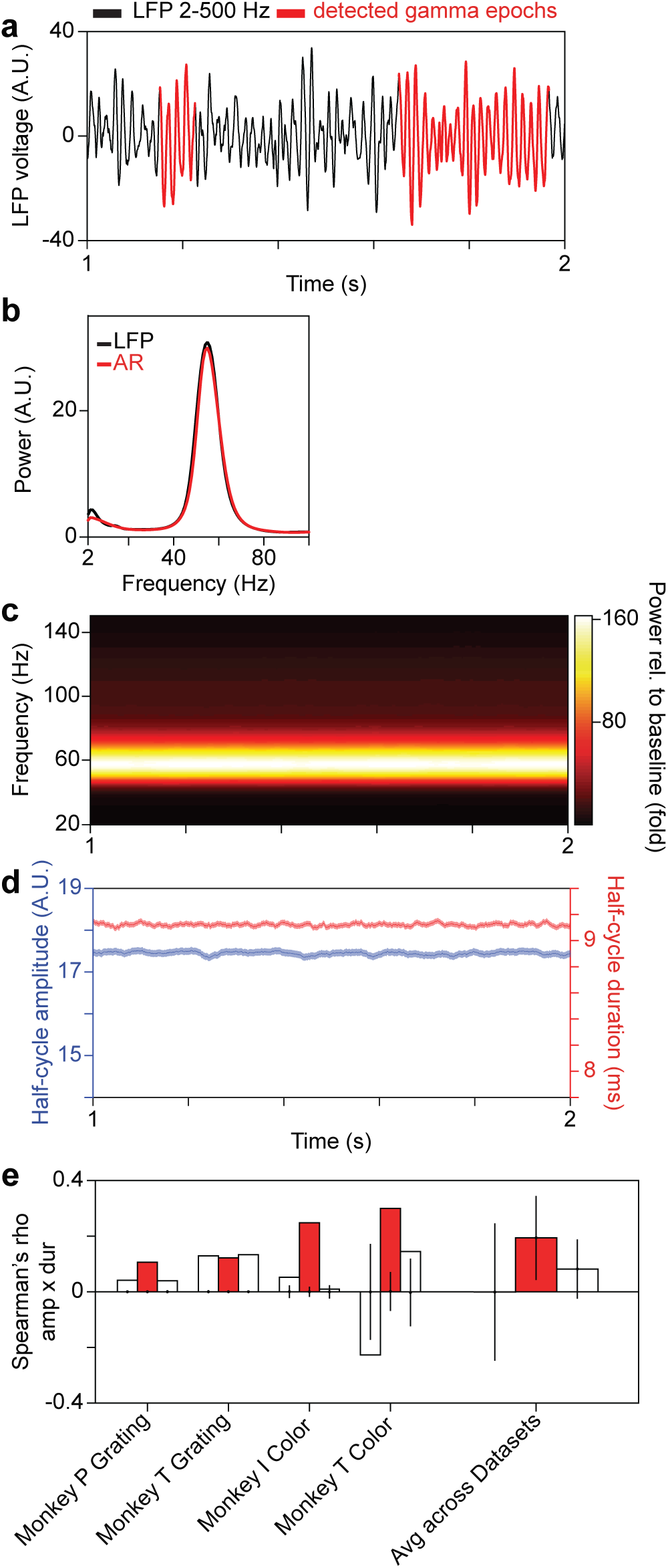
Correlation of gamma-half-cycle amplitudes and durations in an AR model of the visual stimulation period. Panels (**a**-**d**) are based on signals generated by an autoregressive (AR) model of the data used in **Fig. 1a-d**, for the visual-stimulation period, averaged over all selected V1 sites. We refer to the synthetic LFP signal generated by the AR model as AR-based LFP. (**a**) Representative AR-based LFP. Regions presented in red correspond to gamma epochs passing the criterion for stationarity. (**b**) Average raw power of the measured (black) and the AR-based LFP (red). (**c**) Time-frequency power of AR-based LFP. Note the expected absence of temporal trends. (**d**) Time-course of gamma-half-cycle amplitude (blue) and duration (red) of AR-based LFP. Error regions show ±2 SEM based on a bootstrap procedure. (**e**) Same as **Fig. 4a**, but for the AR-based LFP (averages across datasets: red bar: p=0.03, t-test across datasets; white bars p=0.98 and p=0.2, respectively for preceding and succeeding cycles, t-test across datasets).

**Supplementary Figure 4.**
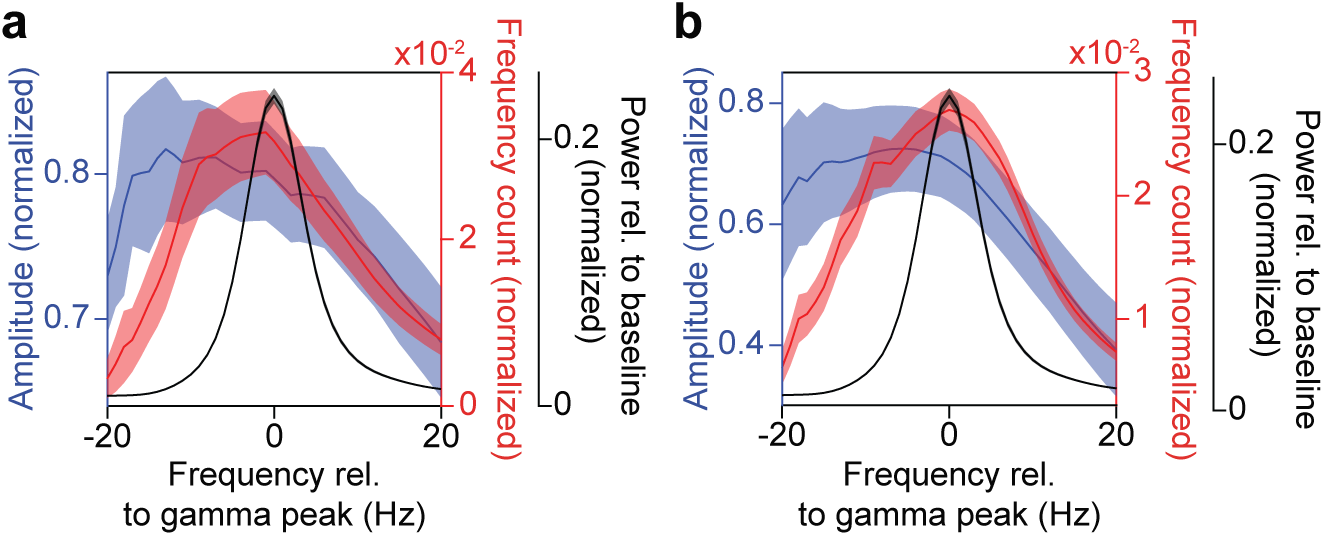
Cycle-based spectra of amplitudes and rates of incidence. (**a**) Same as **Fig. 5a**, and (**b**) same as **Fig. 5b**, but after aligning to the gamma peak in the power-change spectrum.

**Supplementary Figure 5.**
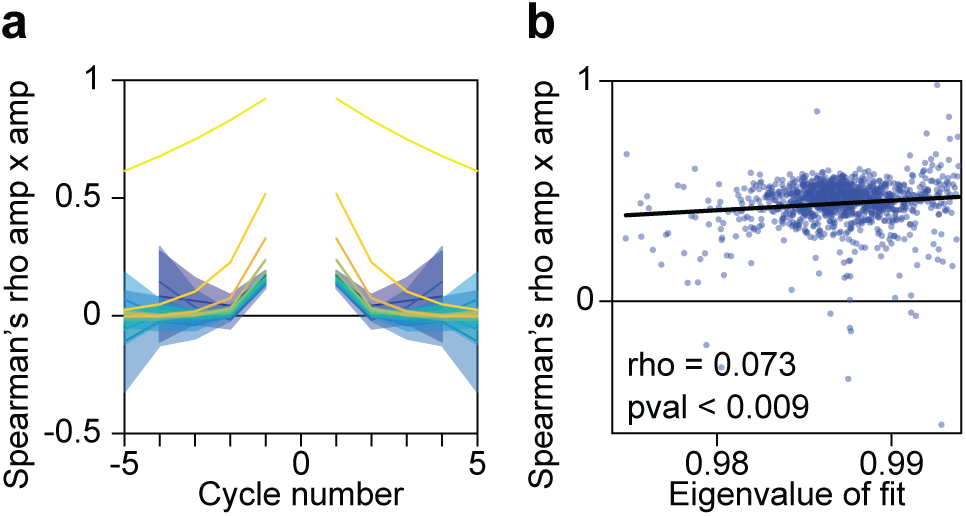
A linear harmonic oscillator driven by noise reproduces LFP gamma-cycle amplitude autocorrelations estimated for full-cycles. (**a**,**b**) Same as, respectively, **Fig. 9a** and **Fig. 9c**, but for full gamma cycles.

**Supplementary Figure 6.**
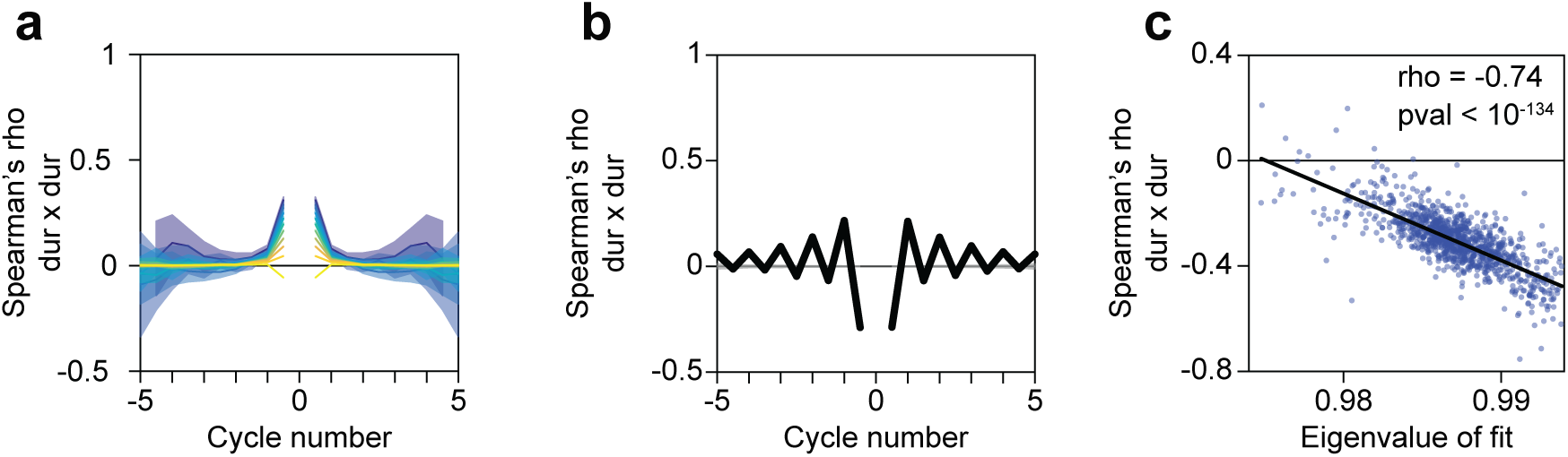
A linear harmonic oscillator driven by noise reproduces LFP gamma-cycle duration autocorrelations estimated for half-cycles. (**a**-**c**) Same as **Fig. 9d-f**, but for gamma half-cycles.

